# Cas13d-mediated isoform-specific RNA knockdown with a unified computational and experimental toolbox

**DOI:** 10.1101/2023.09.12.557474

**Authors:** Megan D. Schertzer, Andrew Stirn, Keren Isaev, Laura Pereira, Anjali Das, Claire Harbison, Stella H. Park, Hans-Hermann Wessels, Neville E. Sanjana, David A. Knowles

**Author notes:** Equal contribution (Andrew Stirn, Keren Isaev).

## Abstract

Alternative splicing is an essential mechanism for diversifying proteins, in which mature RNA isoforms produce proteins with potentially distinct functions. Two major challenges in characterizing the cellular function of isoforms are the lack of experimental methods to specifically and efficiently modulate isoform expression and computational tools for complex experimental design. To address these gaps, we developed and methodically tested a strategy which pairs the RNA-targeting CRISPR/Cas13d system with guide RNAs that span exon-exon junctions in the mature RNA. We performed a high-throughput essentiality screen, quantitative RT-PCR assays, and PacBio long read sequencing to affirm our ability to specifically target and robustly knockdown individual RNA isoforms. In parallel, we provide computational tools for experimental design and screen analysis. Considering all possible splice junctions annotated in GENCODE for multi-isoform genes and our gRNA efficacy predictions, we estimate that our junction-centric strategy can uniquely target up to 89% of human RNA isoforms, including 50,066 protein-coding and 11,415 lncRNA isoforms. Importantly, this specificity spans all splicing and transcriptional events, including exon skipping and inclusion, alternative 5’ and 3’ splice sites, and alternative starts and ends.

## Introduction

Alternative splicing (AS) is a complex, but fundamental, cellular process that regulates approximately 95% of human pre-messenger RNA (pre-mRNA) transcripts (Pan et al. 2008; E. T. Wang et al. 2008; Stanley and Abdel-Wahab 2022). While some AS functions to regulate gene dosage (Lewis, Green, and Brenner 2003; Yan et al. 2015), many splicing products are translated into different protein isoforms, each of which could differ in their cellular localization, protein stability, co-factor or nucleic acid binding, and translation efficiency (Rogalska, Vivori, and Valcárcel 2023). It is well established that AS events are dynamic throughout development and differentiation, especially in brain and muscle tissues (Vuong, Black, and Zheng 2016; Pan et al. 2008; E. T. Wang et al. 2008; Baralle and Giudice 2017). Furthermore, splice altering *cis* genetic variants in a single gene can be sufficient to drive disease, while variants in *trans*-acting RNA binding proteins can disrupt entire splicing networks (Scotti and Swanson 2015; Sisi Chen, Benbarche, and Abdel-Wahab 2021; Seiler et al. 2018). Splicing quantitative trait locus (QTL) analysis has demonstrated that a substantial proportion of complex disease heritability is mediated through genetic effects on AS (Raj et al. 2018; Y. I. Li et al. 2017, 2016; Gusev et al. 2018; GTEx Consortium 2020). Widespread alterations in AS are consistently detected across cancer types, with an estimated 30% more AS events in cancer cells relative to normal cells (Kahles et al. 2018; Stanley and Abdel-Wahab 2022; Danan-Gotthold et al. 2015).

While the importance of AS in human health and disease is well recognized, the functional impact of most AS is unknown, despite the steadily increasing number of annotated isoforms. In the past 5 years alone, the GENCODE annotation has been augmented by approximately 55,000 isoforms for a current total of around 250,000 (Frankish et al. 2021). Therefore, there is a pressing need for methodology to attribute functional roles to specific isoforms.

Historically, the main strategies to identify function at the gene-level has been to knock out, knockdown, or overexpress the gene of interest, but this is more challenging at the isoform-level. Over the last decade, both the experimental and computational methodology to screen for the effects of such perturbations genome-wide have matured, empowered in particular by CRISPR/Cas9 systems. Such CRISPR-based approaches have recently been extended to screen AS events. For example, DNA-based perturbations have been used to delete cassette exons (Thomas et al. 2020; Gonatopoulos-Pournatzis et al. 2020), inhibit transcription from alternative promoters (Davies et al. 2021), or base edit splice sites (Gapinske et al. 2018). RNA-based methods have included RNAi (Celotto and Graveley 2002; Prinos et al. 2011) and antisense oligonucleotides (ASOs) (Villemaire et al. 2003) targeted to unique exonic regions of isoforms, mostly cassette exons. However, these methods have limitations. Mainly, each of these strategies targets a single event type, often focused on exon skipping for cassette exon events. In addition, experimental design is challenging and requires significant computational work to confidently identify isoform expression in a specific cell type and to link a splicing event with its corresponding isoform(s). Conversely, overexpression experiments at the isoform-level, while informative, are low throughput due to the high cost of scaling (Belluti et al. 2021; Gueroussov et al. 2015; Legut et al. 2022).

In this study, we address the needs for both a versatile experimental platform to study isoform-specific function and companion computational tools to simplify experimental design. Our strategy pairs the CRISPR/Cas13d RNA-targeting system with guide RNAs (gRNAs) that target mature RNA exon-exon junctions for specific RNA knockdown. Importantly, this strategy allows targeting of both alternative splicing events, e.g. exon skipping and inclusion and alternative 5’ and 3’ splice sites (SS), and alternative transcriptional events, e.g. alternative starts and ends. While the CRISPR/Cas13 RNA-targeting system has been used to target exonic sequences for efficient RNA knockdown (East-Seletsky et al. 2016; Abudayyeh et al. 2016; Wei et al. 2021; Cheng et al. 2023; Wessels et al. 2023, 2020), it has not been systematically evaluated whether gRNAs that overlap junctions in the mature RNA are similarly effective, as well as specific. To show the broad feasibility of our strategy, we performed the largest junction-targeting Cas13 essentiality screen to date, with 50,512 gRNAs targeting 6,957 junctions across 837 essential genes.

To optimally analyze the screen data, we developed a novel Bayesian linear mixed model, SEABASS (Screen Efficacy Analysis with BAyesian StatisticS), which robustly integrates multiple time-points and replicates. We then compared adaptations of our TIGER deep learning model (Wessels et al. 2023) to investigate which sequence and non-sequence features contribute to junction-level versus exon-level gRNA efficiency. We find that rules governing Cas13d gRNA efficacy are similar for both exons and junctions. Given the complexity of experimental design, we additionally developed an R package, Isoviz, which integrates transcript structures from GENCODE or long read sequencing data, spliced read counts per junction, and TIGER gRNA efficiency predictions. We highlight the usefulness of Isoviz and validate TIGER predictions on multiple genes using both quantitative reverse transcription PCR (RT-qPCR) and long read RNA-sequencing analyses. We confirm that TIGER predictions strongly correlate with measured RNA knockdown and show that junction-specific knockdown corresponds to the expected isoform-specific knockdown. Our unified experimental and computational approach represents the most versatile and efficient platform for isoform-specific knockdown to date.

## Results

### Cas13d essentiality screen shows broad applicability of junction-targeting strategy

Our proposed CRISPR/Cas13 strategy to utilize gRNAs at exon-exon junctions is a versatile method that aims to target unique sequences of alternative start and end exons, alternative 5’ and 3’ SS, and cassette exons without inadvertently cutting the pre-mRNA. To more easily relate this junction-level approach with isoform-level outcomes, we categorized junctions into three types: (1) common across all isoforms of a gene (“common”), (2) exclusive to one isoform (“fully unique”), and (3) shared across a subset of isoforms (“partially unique”) (Figure 1A). Using this junction classification, we evaluated the potential of our junction-centric strategy to functionally interrogate transcript isoforms. Across human GENCODE v41 (Basic Annotations, containing 18,568 protein-coding and 14,849 lncRNA genes) (Frankish et al. 2021) 53,890 protein-coding and 23,502 lncRNA junctions are uniquely targetable, corresponding to 36,604 and 11,053 isoforms, respectively (“Fully Unique” in Figure 1B; Mouse vM33 in Supplemental Figure 1A). Additionally, isoform expression is often tissue-specific, so that partially unique GENCODE junctions may be re-classified as fully unique within specific cell types. By combining fully and partially unique categories, we estimated that our junction-centric strategy has the potential to target upwards of 97% of protein-coding and lncRNA isoforms from multi-isoform genes, respectively, highlighting its broad potential for isoform-specific functional studies.

**Figure 1.**
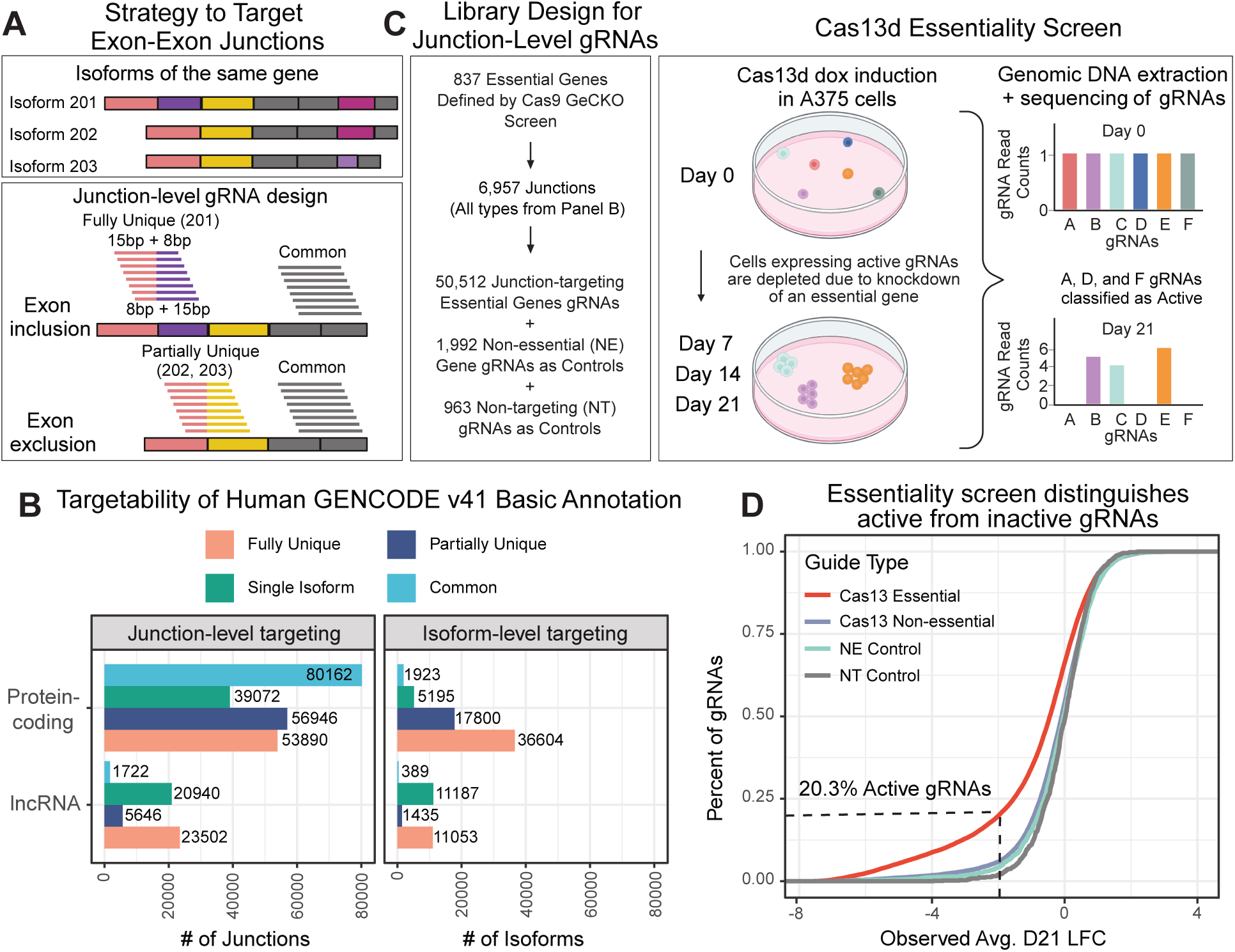
Cas13d essentiality screen shows broad applicability of junction-targeting strategy. (A) The exon-exon junction-targeting strategy and junction categories relating junction-level information to isoform-level information for a hypothetical gene. A common junction (gray exons) is shared across all isoforms, a fully unique junction (pink/purple exons) is specific to the 201 isoform, and a partially unique junction (pink/yellow exons) is present in a subset of isoforms, 202 and 203. For each junction, eight 23bp gRNAs are considered, with a starting position 15bp upstream of the junction and tiling every base pair to starting position of 8bp upstream of the junction. (B) Counts of theoretically targetable junctions and isoforms in GENCODE v41 (Basic Annotations, containing 18,568 protein-coding and 14,849 lncRNA genes), broken down by categories described in (A). (C) Left: the number of essential genes and their respective junctions that are targeted in the screen design. Right: the logic of the essentiality screen, where cells expressing gRNAs that knockdown essential genes are depleted over the 21 day time course. (D) Cumulative density function plot comparing Day 21 LFC between the re-defined Cas13d essential gene (n=27,804 gRNAs across 481 genes define by requiring more than 5% active gRNAs/gene) and Cas13d non-essential gene gRNAs (n=22,708 gRNAs across 356 genes) plus the two sets of control gRNAs for non-essential (NE, n=1,992 gRNAs) genes and non-targeting (NT, n=963 gRNAs). The dotted black lines mark the cutoff for active gRNAs defined as having LFC <0.01 quantile of NT gRNA normal distribution.

Motivated by this potential widespread transcriptome coverage, we designed an unbiased essentiality screen to study whether Cas13d-gRNA complexes could efficiently access their target RNA splice junctions at genome-scale. Complex-target formation could be sterically hindered by the exon-junction complex (Hir, Saulière, and Wang 2015) or splicing factor binding. Our screen contains the largest number of junction-targeting gRNAs to date, with 50,512 gRNAs across 6,957 junctions from the top 837 essential genes, defined by the Cas9-based GeCKO screen in A375 cells (Shalem et al. 2014). Importantly, we selected 23bp gRNAs tiling each junction from -15bp to +15bp, expecting a mix of active and inactive gRNAs, so that each junction is represented by eight gRNAs shifted by 1bp. We include gRNAs for all types of junctions classified in Figure 1B, with the majority targeting single isoform and common junctions to give total gene knockdown (Supplemental Figure 1B). In addition, we included two sets of control gRNAs: 1,992 non-essential (NE) gene and 963 non-targeting (NT) gRNAs (Figure 1C). The pool of lentivirus gRNAs were transduced in two biological replicates into monoclonal, single-integration Cas13d-expressing A375 cells. We quantified gRNA depletion at Days 7, 14, and 21 after Cas13d induction relative to Day 0 using log2 fold changes (LFC) of gRNA read counts. We used gRNA depletion as a surrogate for RNA knockdown, i.e., a gRNA with a significant negative LFC corresponds to an active gRNA that effectively knocks down its essential gene target (Figure 1C). We observed gRNA depletion at D21 relative to NT controls across all junction types (Supplementary Figure 1B).

Different screening platforms are known to return non-overlapping sets of essential genes, especially when comparing gene knockout versus knockdown (Morgens et al. 2016; Sanson et al. 2018). We designed our screen using essential genes identified through Cas9 gene knockout in GeCKO (a DNA-targeting system), expecting that some genes would not be essential in our RNA-targeting system, where the gene is only knocked down. This discrepancy could bias gRNA efficacy estimation if a gene’s gRNAs are mis-classified as inactive only because the knockdown was not sufficient to reduce cell proliferation. To avoid this bias we reclassified genes with less than 5% active gRNAs, i.e. fewer than 1 gRNA depleted out of 20 (LFC < -1.96, corresponding to the 0.01 quantile of the NT gRNA’s normal distribution) as being non-essential in our screen (termed “Cas13 Non-Essential”). This removed 356 genes from future analyses. Of the remaining 481 Cas13d essential genes, 20.3% of their 27,804 gRNAs are active (Figure 1D). This percentage is similar to active gRNA estimates from other recent Cas13d unbiased essentiality screens (Wei et al. 2021; Cheng et al. 2023; Wessels et al. 2023).

When evaluating the quality of our screen data, we observed strong concordance in essentiality scores for shared genes between our screen and previous DNA and RNA based essentiality screens in A375 cells (R = 0.47-0.64 for 1,165 genes; Supplementary Figure 1C) (Doench et al. 2016; Sanson et al. 2018; Shalem et al. 2014; Tsherniak et al. 2017). This included strong correlations of scores across replicates and time points within our screen (R = 0.76-0.92, Supplementary Figure 1C). Additionally, we examined 258 gRNAs shared between our A375 screen and a previously published HEK293FT screen; we noted strong correlation of LFCs across our two biological replicates in A375 and between our A375 results and the HEK293 scores (Supplementary Figures 1D; (Wessels et al. 2023)). Finally, to mitigate concerns of potential Cas13d collateral activity (Q. Wang et al. 2019; Ai, Liang, and Wilusz 2022; Shi et al. 2023), we examined gRNA depletion of the 1,992 NE gene control gRNAs (413 genes ranging from 10-492 transcripts per million (TPM)). Reassuringly, the knockdown of these non-essential genes by Cas13d did not lead to cell death and thus, gRNA depletion in our screen (Supplemental Figure 1E). These analyses affirm that our Cas13d junction-targeting strategy is feasible and broadly capable of knocking down RNA while offering the distinct advantage of isoform-specific knockdown.

### Deep learning uncovers similar rules for gRNAs targeting junctions and exons

The results from our junction-targeting essentiality screen indicated that a substantial proportion of junction-spanning gRNAs can effectively knockdown RNA. However, we recognized that junctions are different from exons. They are bound by different proteins and have limited availability in the nucleus for Cas13/gRNA binding depending on splicing efficiency, position in the transcript, and time to export. Junction-targeting may therefore merit different considerations when designing efficient gRNAs. Current models to predict Cas13d gRNA efficiency, including our previously published TIGER deep learning model, were trained primarily on exon-targeting gRNAs (Wessels et al. 2020; Wei et al. 2021; Cheng et al. 2023; Wessels et al. 2023). It is unclear whether a deep learning model would learn additional (or different) features relevant to junction-targeting if trained on junction gRNA data. As our screen includes more junction-targeting gRNAs than any previous Cas13 screen, we are well-positioned to investigate this question.

To obtain the cleanest possible input for model training, we wanted to utilize data from all time points and replicates in our screen. We developed SEABASS, a robust Bayesian linear mixed model (see Methods) that takes as input LFCs for D7, D14, and D21 across both biological replicates and outputs a slope (LFC over time) for each gRNA (Figure 2A). A SEABASS negative slope indicates a more active gRNA, while a slope closer to 0 indicates an inactive gRNA. We retrained several adaptations of our previously published TIGER deep learning model (Wessels et al. 2023) available at tiger.nygenome.org. The table in Figure 2B details the original TIGER model trained on exon LFC data, TIGERjunction, trained on junction LFC data, TIGERbass, trained on junction SEABASS slopes, and TIGERsite, trained on junction LFC data with additional sequence context as input (Figure 2B).

**Figure 2.**
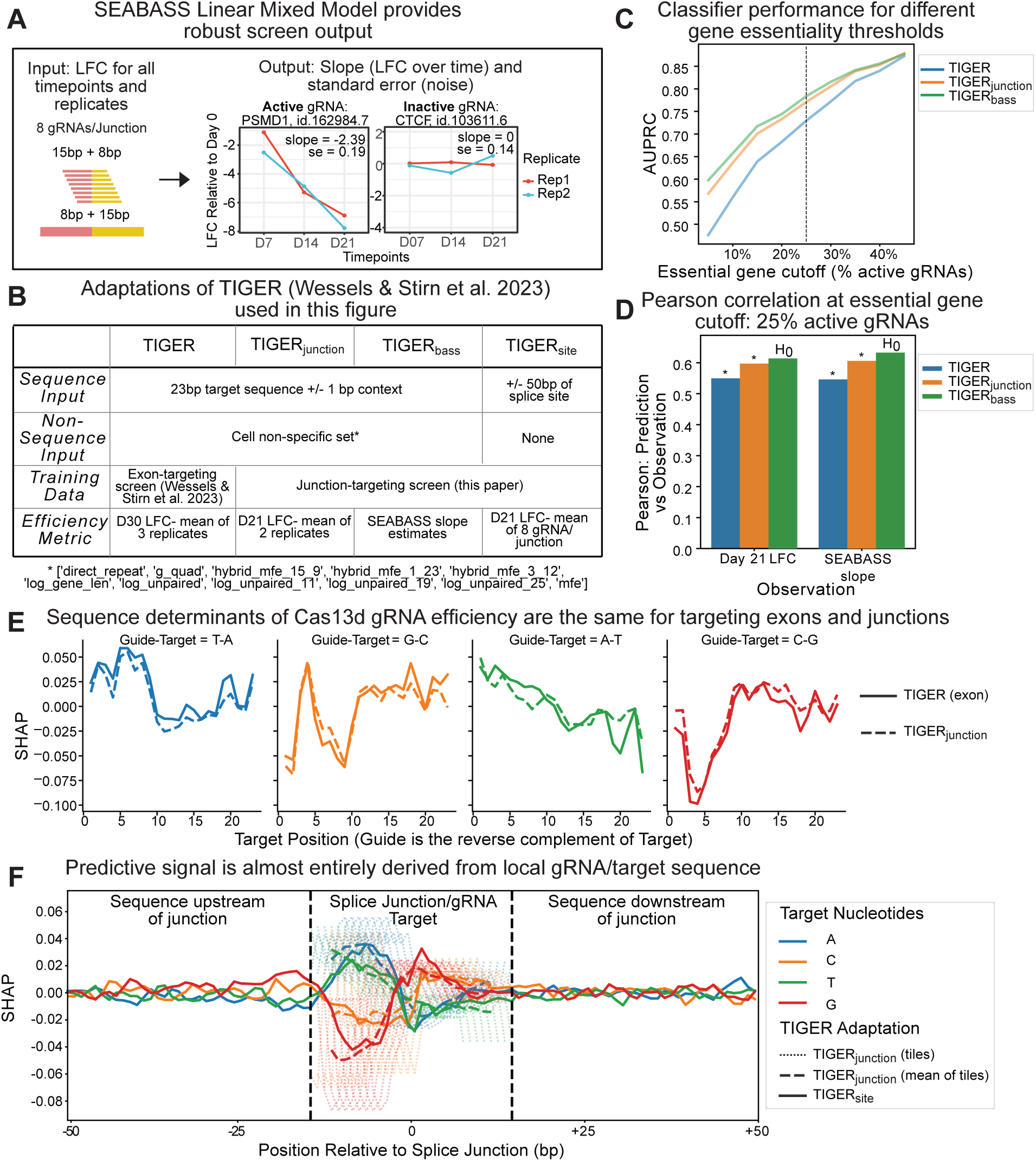
Deep learning uncovers similar rules for gRNAs targeting junctions and exons. (A) Schematic of SEABASS linear mixed model. Inputs are LFCs for all timepoints and replicates for junction-spanning gRNAs in our essentiality screen. The output is a slope, where a more negative slope corresponds to a more active gRNA. (B) Table comparing adaptations of TIGER used in this figure. The non-sequence input list for ‘Cell non-specific set’ is given below the table. (C) Cross-validated (see TIGER Model Architecture, methods) area under the cross-validated precision-recall curve (AUPRC) across different gene essentiality cutoffs. The percentage on the x-axis is the proportion of required active gRNAs per gene to define a gene as essential. The dotted line corresponds to 25% which is the cutoff used in D. (D) Cross-validated Pearson correlations between predicted and observed screen data at the 25% gene essentiality cutoff. X-axis distinguishes which type of observed data (LFC or slopes) was used in the correlations. *p<0.0001. (E) SHAP plots comparing sequence preferences for TIGER (solid line) and TIGER_junction_ (dashed line). The line colors are based on target nucleotide (e.g. A is blue), which is given at the top, along with the corresponding gRNA nucleotide, which is the reverse complement. A more negative SHAP value indicates the model will output a lower LFC estimate (i.e. predict stronger Cas13 activity) when that position has the specific nucleotide. (F) SHAP values from two adaptations of TIGER. TIGER_site_ (solid lines) predicts the average LFC of the eight tiled gRNAs in our screen from the full junction sequence (-50bp to +50bp about the splice junction). TIGER_junction_ predicts a gRNA’s LFC from the 23bp target sequence. We tile the SHAP values from TIGER_junction_ across the junction at the same eight positions as the gRNAs in our screen (tiles; dotted lines). Vertical gray bars mark the range of gRNA positions overlapping the splice site. We positionally average (ignoring positions with less than three overlapping tiles) the tiled SHAP values from TIGER_junction_ (tile mean; dashed lines) and find that it closely recapitulates the SHAP values from TIGER_site_. The x-axis is the gRNA target position centered at the splice site.

To evaluate the classification of active versus inactive junction-targeting gRNAs by TIGER, TIGERjunction, and TIGERbass, we generated cross-validated precision-recall (PR) curves, using observed screen data as labels. Using increasingly more stringent gene essentiality cutoffs for the observed test data (5-45% active gRNAs/gene), we compared the area under precision-recall curves (AUPRC) across the three TIGER adaptations (Figure 2C). As we filter for genes with a higher proportion of active gRNAs, we have more confidence that these genes are truly essential and that the observed active/inactive gRNA classifications are correct. TIGERbass performs the best at a lower cutoff, when the data is noisier, but all three models converge at the most stringent essentiality cutoff (45%) to an AUPRC of 0.88 (Figure 2C). At a more modest cutoff of 25%, where we have more gRNAs and genes for model assessment, we observed similar trends in Pearson correlation values between predictions and observed screen data (Figure 2D). TIGERbass not only outperforms the other models when tested against its predictive modality of SEABASS slopes but also outperforms TIGER and TIGERjunction at their predictive modality of LFC, confirming the benefit of SEABASS’s de-noising of the screen data (R = 0.63 for TIGERbass versus R = 0.61 and 0.55 for TIGERjunction and TIGER, respectively), although by a small margin (Figure 2D).

Following previous work (Wessels et al. 2023; Wei et al. 2021), we employed the model interpretation technique SHapley Additive exPlanations (SHAP) (Lundberg and Lee 2017) to assess features used by the model to predict gRNA efficacy. First, we focused on nucleotide preferences for active gRNAs by comparing SHAP values across the 23bp target RNA sequence for TIGER, which is trained on exon LFC data alone (Wessels et al. 2023) versus TIGERjunction, which is trained on junction LFC data alone (dashed line; Figure 2E). Pearson correlations of positional nucleotides and observed LFC data produced similar plots. (Supplemental Figure 2A). The strong agreement of both the SHAP and Pearson plots between an exonic dataset and our junction dataset suggest that sequence determinants of Cas13d gRNA efficiency are indistinguishable for targeting junctions and exons.

Next, we asked whether there was broader sequence context surrounding junctions that influence gRNA efficiency, for example as a result of secondary structure or sequence-specific splicing factor binding. We compared two adaptations of TIGER: 1) TIGERjunction that is trained using 23bp of target RNA sequence and 2) TIGERsite that considers -50/+50bp around each junction site (Figure 2B). We trained TIGERjunction to predict a gRNA’s LFC and trained TIGERsite to predict the mean of the eight gRNAs’ LFCs that target each junction. When we tiled TIGERjunction SHAP values across the junction as we tiled gRNAs across the junction in our screen (TIGERjunction tiles) and positionally averaged TIGERjunction SHAP values (TIGERjunction tile mean), we recovered almost exactly the TIGERsite SHAP profile within the tiling window (between the gray dotted lines; Figure 2F). This suggests that TIGERsite learns to average the sequence preferences of the eight possible gRNA alignments despite having the opportunity to learn novel junction sequence preferences. We conclude from this analysis that the predictive signal is almost entirely derived from the local sequence of the gRNA/target, with minimal if any contribution from broader sequence context. This is in line with findings from Wessels et al. (2023).

TIGERbass input included 23bp of target sequence and a set of cell non-specific features relating to RNA structure (Figure 2B). From this, the model achieved a Pearson correlation of 0.59 between predicted and observed slopes (5% gene essentiality cutoff; Supplemental Figure S2B). To explain discrepancies between the predicted and observed results, we calculated residuals for each gRNA and investigated their association with a set of non-sequence features, not included in TIGERbass training (Supplemental Figures S2C). We considered RNA half-life estimates, gene expression, and nuclear RNA localization in A375 (Cas13d is localized to the nucleus), gene length, intron length, relative junction position in a gene, and gRNA tiling position. We fit a linear regression to evaluate feature associations with residuals, but found negligible relationships across all non-sequence features tested (Supplemental Figures 2D-E). To further support these findings, we found that SHAP values from non-sequence features matched those for a model trained on exonic target sites only (data not shown). We conclude that gRNA sequence, and thus its target RNA sequence, are the primary features predictive of Cas13d gRNA junction-targeting efficiency. We hypothesize that this will hold true for future RNA-targeting CRISPR systems. Practically, this means that one can use existing prediction tools for junction-targeting gRNA design such as TIGER or other models (Wei et al. 2021; Cheng et al. 2023).

### Isoviz automates experimental design for Cas13d-based isoform-specific knockdown

We have shown that mature RNA exon-exon junctions are broadly targetable using Cas13d in a screen platform and that existing gRNA prediction tools can be used to select effective gRNAs that span these junctions. However, given the complexity of AS in mammalian cells, it is significantly more challenging to design and interpret results for Cas13-based experiments that target junctions and their isoforms compared to those targeting exons. First, there is uncertainty regarding which isoforms are expressed and at what levels in a given cell type. Furthermore, some AS events, especially alternative 5’ and 3’ SS, are difficult to visualize from a typical transcript diagram but could have substantial consequences, especially if they introduce a frameshift. Finally, selecting efficient gRNAs that span specific junctions can be time-consuming. The individual resources and tools currently available to address these challenges are disjointed and require extensive manual, error-prone labor.

To aid in designing isoform-centric experiments, especially CRISPR-based ones, we developed Isoviz, an R package that integrates transcript-level annotations from GENCODE or long read analyses, spliced read counts for each junction, and TIGER gRNA predictions (Figure 3A). The result of this integration shows a clear link between each junction, its expression within a cell type, and its corresponding isoform(s). Based on this, a user can quickly design gRNAs to junctions that map uniquely or partially uniquely to knockdown an isoform of interest.

**Figure 3.**
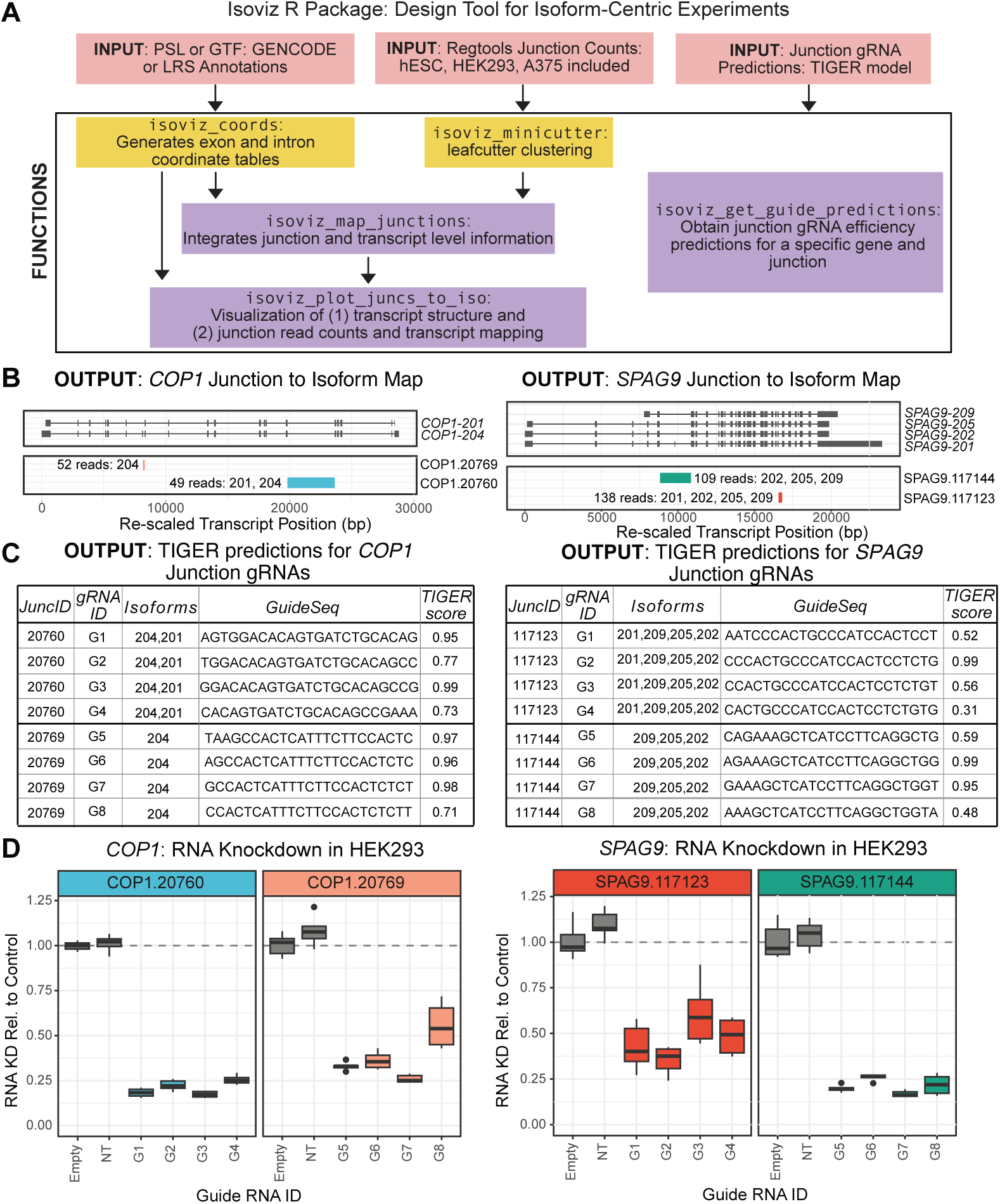
Isoviz automates experimental design for Cas13d-based isoform-specific knockdown. (A) Isoviz takes as input three files, depending on the function (red boxes). Default input files are already provided on github, but the user can incorporate their own inputs. Pre-processing functions (yellow boxes) reformat input data for downstream plotting functions. Main visualization and table output functions (purple boxes) are run on a per-gene basis, as specified by the user. (B) Example Isoviz visualization for COP1 and SPAG9 isoform structures (top) and two selected junctions per gene plotted as introns (bottom). The user can specify if all or a subset of junctions should be plotted. The text next to each intron corresponds to the number of reads overlapping that junction and the specific isoforms containing that junction. (C) Example Isoviz output table showing gRNAs targeting junctions in (B) and their corresponding TIGER scores. (D) COP1 and SPAG9 RNA knockdown using gRNAs from (C) to target junctions in (B). RNA knockdown is measured by RT-qPCR relative to cells transfected with an empty gRNA cassette as a control (4 guides tested per junction). Junction IDs (20760 and 20769, 117123 and 117144) and boxplot colors correspond to junctions in (B). Each boxplot represents three RT-qPCR measurements from two biological replicates.

We first illustrate the utility of Isoviz in the design of junction-based RNA knockdown experiments for two genes with relatively simple transcript structures, *COP1* and *SPAG9*. Isoviz allows for the visualization of all junctions and their corresponding isoforms, followed by a table with the top TIGER scoring gRNAs for each junction, with the option to filter for a subset of junctions, as in Figures 3B-C. Figure 3B identifies that the COP1.20769 junction is specific to the *COP1-204* isoform, and SPAG9.117144 is partially unique and maps to all isoforms except *SPAG9-201* (junction IDs are assigned in Isoviz for ease of comparison). To select gRNAs that target these junctions, Figure 3C shows four of the eight possible gRNA options per junction and their TIGER score. Both visualization and table outputs save the user significant time and reduce the possibility of error.

Next, we wanted to test COP1 and SPAG9 junction-specific gRNAs that were designed using Isoviz and directly measure the levels of RNA knockdown. We created a non-viral, piggyBac-Cas13 vector system by cloning a doxycycline (dox)-inducible Cas13d (NLS-*Rfx*Cas13d-NLS) and a gRNA cassette (hU6-DR-BsmBI) (Wessels et al. 2020) into two separate piggyBac transposon backbones (Schertzer et al. 2019), creating PB-Cas13 and PB-rtTA-gRNA, respectively (Ding et al. 2005; G. Wang et al. 2017; S. Li et al. 2017). We generated stable HEK293 cell lines for the 16 *COP1* and *SPAG9* gRNAs depicted in Figures 3B-C and measured RNA knockdown, with junction-specific primers, via RT-qPCR after 24 hours of dox induction of Cas13d. All *COP1* and *SPAG9* gRNAs in Figure 3C produced robust and consistent knockdown across two biological replicates, ranging from 39-83% reduction in RNA expression relative to control (Figure 3D, Supplemental 3A). We further show that RNA knockdown for all 8 *COP1* gRNAs was similar in hESCs and across different timepoints of Cas13 dox-induction (Supplemental 3B). Importantly, robust knockdown (>80%) can be observed as early as 12 hours post dox-induction. We use the 24 hour time point for all future experiments. In line with a previous study (Burris et al. 2022), we found that RT-qPCR primers flanking or overlapping the gRNA cutsite showed the most RNA knockdown, and we designed all RT-qPCR primers for future experiments accordingly (Supplemental Figure 3C). Collectively, our Isoviz design tool and robust technical optimization provide a powerful framework for others to effectively execute junction and isoform centric experiments.

### PacBio long read sequencing confirms isoform-specific knockdown using junction gRNAs

We have shown that junction-targeting gRNAs achieve consistent and robust RNA knockdown across diverse cell types, replicates, and timepoints. A crucial next step is to determine how junction-centric RNA knockdown quantification relates to desired isoform-specific knockdown. We chose *RBFOX2* as a case-study because of its high isoform complexity. To accurately identify which *RBFOX2* isoforms are expressed in hESCs, we performed PacBio long read RNA-sequencing (LRS) (Al’Khafaji et al. 2023). Importantly, for all LRS analysis, we grouped isoforms that have identical sets of splice junctions, thereby excluding differences in transcription start and end sites. Based on the LRS analysis, we identified 12 *RBFOX2* isoforms expressed in the unperturbed hESC control, six of which have not been previously annotated in GENCODE (Figure 4A, top panel; isoforms > 0.6 TPM or 10 full length reads in the control).

**Figure 4:**
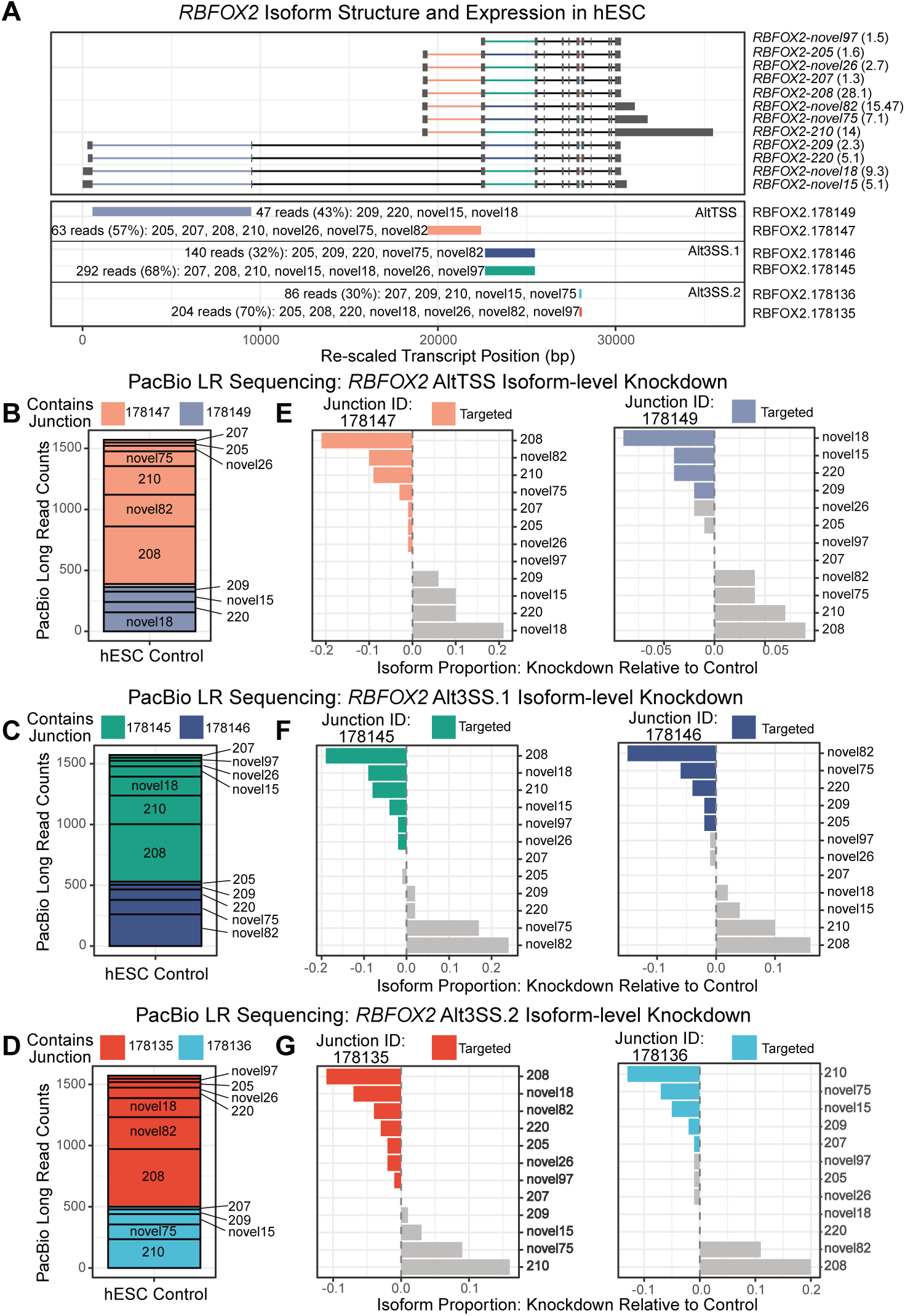
PacBio long read sequencing confirms isoform-specific knockdown using junction gRNAs. (A) Isoviz plot of all annotated and unannotated RBFOX2 transcripts detected by PacBio long read sequencing in hESCs (top; isoforms > 0.6 TPM or 10 full length reads in the control). TPM in hESC control for each isoform is in parentheses to the right of the plot, next to the isoform name. The bottom panel shows six selected junctions plotted as introns from three different events: an alternative TSS (AltTSS), an alternative 3’ SS differing by 3bp (Alt3SS.1), and an alternative 3’ SS differing by 12bp (Alt3SS.2). The text next to each intron gives the number of junction-spanning short reads from Illumina sequencing, along with the percentage of reads mapping to that junction versus its alternative junction. Additional text identifies transcripts that contain that specific junction. (B-D) Stacked bar plots show the proportion of PacBio full length reads mapping to each RBFOX2 transcript in the hESC control, arranged and colored by whether they contain a specific junction. (E-G) Barplots showing change in transcript proportion upon knockdown with the indicated gRNA relative to the control for the two alternative junctions for the AltTSS (E), Alt3SS.1 (F), and Alt3SS.2 (G) that were depicted in (A). Each bar represents a transcript and is colored according to whether it contains the junction being targeted for knockdown (color matches intron color in A) or it does not (gray).

To inform our *RBFOX2* experimental design, we used Isoviz to integrate LRS transcript-level information with matched short-read junction-level counts. The complex transcript structure of *RBFOX2* harbors three alternative TSS (AltTSS), a cassette exon (not shown), and two separate alternative 3’ SS events, where the two junctions at Alt3SS.1 differ by 3bp and the two junctions at Alt3SS.2 differ by 12bp (Figure 4A, bottom panel). The subtle base pair differences at Alt3SS.1 and Alt3SS.2 junctions cannot be distinguished by eye, exemplifying another way that Isoviz visualization is helpful for accurately identifying small splicing changes and mapping them to their corresponding isoform(s). We did not test junction gRNAs at the cassette exon event due to low expression of the exclusion product present in *RBFOX2-205*, *RBFOX2-207*, and *RBFOX2-novel26*, with TPMs of 1.6, 1.3, and 2.7, respectively (each < 2% of all *RBFOX2* reads; Figure 4A). To test the specificity of RNA knockdown in hESCs using junction gRNAs, we used Isoviz and TIGER predictions to select one gRNA for each of the six junctions in Figure 4A.

We confirmed RNA knockdown using two complementary but distinct methods: RT-qPCR to show junction-specific knockdown and LRS to confirm isoform-specific knockdown. Junction-specific RT-qPCR in hESCs shows RNA knockdown of the gRNA-targeted junction, with minimal effects on the alternative junction (Supplemental Figure 4A). We obtained similar results in HEK293 cell lines (Supplemental Figure 4B). Using LRS data to evaluate isoform-specific knockdown, we first aggregated isoform expression by junction for each splicing event (AltTSS, Alt3SS.1, Alt3SS.2) in the unperturbed hESC control (relative proportions in stacked barplots; Figures 4B-D). Figures 4E-G illustrate the expected and dramatic shift in *RBFOX2* isoform ratios that occurred upon knockdown. Across all six junction-targeting knockdown experiments, we observed the expected depletion or enrichment of specific *RBFOX2* isoforms based on whether they contain the targeted junction or not, respectively (Figures 4E-G, Supplemental Figure 4C). For example, *RBFOX2-208* is the isoform with the highest proportion of reads in the control and contains junctions 178147, 178145, and 178135 (Figure 4B-D). Thus, when any of these three junctions are targeted by a gRNA, the proportion of *RBFOX2-208*, relative to all other *RBFOX2* isoforms, decreases the most, between 10-20% (Figures 4E-G, barplots on the left). This corresponds to a reduction of 13-25 TPM, a ∼89% knockdown of its total expression (Supplemental Figure 4C). Conversely, when *RBFOX2-208* is not targeted by each of the other three gRNAs, its relative proportion increases the most, between 8-20% (Figures 4E-G, barplots on the right). These results confirm that targeting unique or partially unique exon-exon junctions is highly specific in knocking down its intended isoform target(s). Remarkably, this remains true for junctions that differ by only three nucleotides, as in Alt3SS.1 (Figure 4F).

As an additional example of junction-specific knockdown, we designed gRNAs for two unique junctions targeting the alternative last exons of *MKNK2* (Supplemental Figure 4D; (Maimon et al. 2014). In HEK293 cells, we measured RNA knockdown of 78% at the 128207 junction of *MKNK2-202* and 87% at the 128208 junction of *MKNK2-201* (Supplemental Figure 4E). Importantly, the alternate *MKNK2* junction in each experiment was minimally affected. Together, the *RBFOX2* and *MKNK2* experiments emphasize the versatility of our junction strategy to target a broad range of transcriptional and splicing events that can be leveraged to achieve isoform-specific RNA knockdown.

### Measured RNA knockdown validates models trained on essentiality data

Model performance when predicting Cas13d gRNA efficiency has primarily been evaluated using LFCs from essentiality screens (Wessels et al. 2023; Wei et al. 2021; Cheng et al. 2023). While essentiality screens conveniently allow for the testing of thousands of gRNAs at once, they provide only a proxy for the level of RNA knockdown. Ideally, model performance would be evaluated by comparing predictions directly to RNA knockdown measurements.

Here, we aggregated all of the HEK293 RT-qPCR data generated in this paper and compared observed RNA knockdown to predictions from three publicly available deep learning models for Cas13 gRNA efficacy prediction, TIGER (Wessels et al. 2023), Wei et al. (2023), and Cheng et al. (2023). TIGER performs the best (R = -0.86; Figure 5A), with TIGERbass and TIGERjunction models from Figure 2 also showing a strong correlation of -0.78 and -0.81, respectively (data not shown). Additionally, Wei et al. (2023) achieved a good correlation of -0.74 and Cheng et al. (2023) shows a -0.56 correlation (Figure 5A). All models, except Cheng et al., agreed on which gRNAs would have very low activity (upper left corner, Figure 5A). Indeed, these gRNAs did not significantly knockdown RNA. This suggested to us that these models are especially good at calling inactive gRNAs, but vary in their ability to predict the precise level of RNA knockdown that will be achieved with an active gRNA.

**Figure 5:**
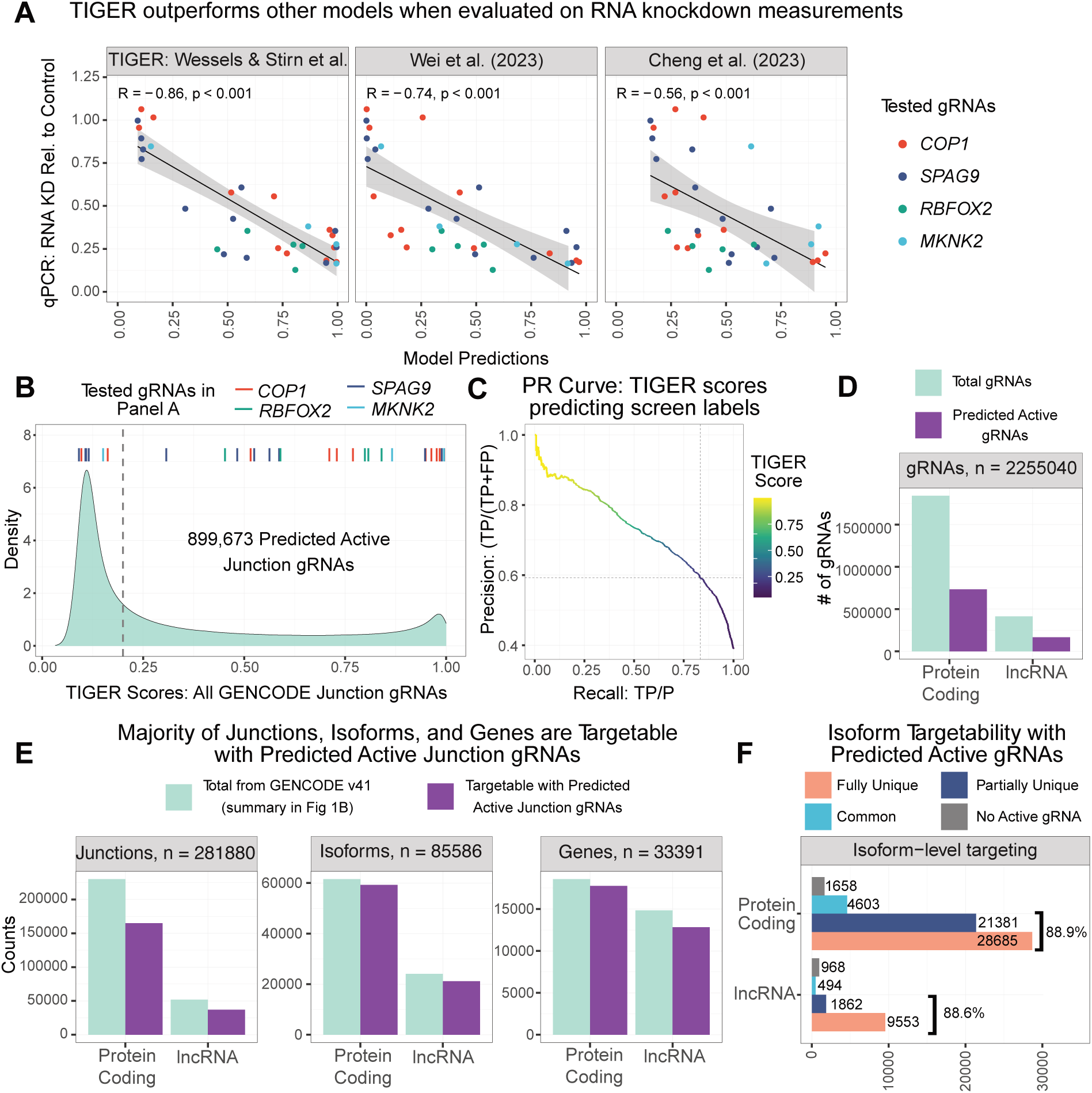
Measured RNA knockdown validates models trained on essentiality data. (A) Pearson correlations between measured RNA knockdown by RT-qPCR and model predictions from TIGER and two additional previously published Cas13 gRNA prediction tools. RT-qPCR data is aggregated from the data in Figures 3 and 4 in HEK293 cells. Each point represents an average of at least two biological replicates with at least two technical replicates. (B) TIGER scores (Wessels et al. 2023) for all GENCODE v41 basic annotation junction-spanning gRNAs with 8 gRNAs tiling each junction from - 15/+8 bp to -8/+15 bp. gRNAs with homopolymers (runs of G’s, C’s, A’s, and T’s) and those mapping to multiple gene loci were excluded. The colored bars at the top mark the scores for each gRNA plotted in (A). Based on predictive scores for gRNAs that knocked down their RNA target in (A), we suggest gRNAs with TIGER scores >0.2 (gray dashed line) are likely active. (C) Precision-Recall Curve assessing TIGER classification of active and inactive junction-targeting gRNAs across varying TIGER score thresholds (0-1). Junction gRNAs from our essentiality screen in Figures 1-2 (filtered for genes with > 25% active gRNAs) were used as labels (active defined as LFC <0.01 quantile of NT gRNA Normal distribution). The gray dotted lines mark the precision and recall values when using the suggested 0.2 TIGER score threshold. (D) The total number of possible junction gRNAs from GENCODE (green) and the number with a TIGER score of greater than 0.2 (purple) for both protein-coding and lncRNA genes. (E) The total number of junctions, isoforms, and genes annotated in GENCODE (green, ‘n’ value at the top of each plot) versus the number that are likely targetable with active junction-spanning gRNAs (score > 0.2, purple). (F) Adjusted counts of isoform-level categories from Figure 1B, considering targetability with predicted active gRNAs (isoforms with no active junction-spanning gRNAs are in gray). We excluded the 5,195 protein-coding and 11,187 lncRNA single isoform genes so these would not be used in isoform-specific functional studies.

Motivated by TIGER’s strong performance in predicting junction-level RNA knockdown, we used TIGER to score eight tiling gRNAs per junction across all GENCODE human basic annotation transcript junctions (1.8 million gRNAs after filtering out gRNAs with homopolymers and non-unique sequence; Figure 5B). To set a score threshold to distinguish active versus inactive gRNAs, we considered two sets of data. First, we used our observed RNA knockdown data from RT-qPCR (see ticks at top of Figure 5B). Second, we assessed precision and recall across varying TIGER score thresholds to classify active and inactive gRNAs from our essentiality screen (Figure 5C). Based on these data, we suggest a threshold of 0.2 (gray dashed lines in Figures 2B-C) which accurately separates active and inactive gRNAs in our RT-qPCR measurements and gives a precision of 0.59 and a recall of 0.83 on our screen data. However, this threshold is flexible and can be selected by the user based on their needs. For example, when designing a future screen, a higher precision may be desirable to ensure most gRNAs are active.

Using the 0.2 threshold, 899,673 junction-spanning gRNAs are predicted to be active (Figure 5D). These active gRNAs target 201,905 junctions across 80,440 isoforms, belonging to 17,752 protein-coding and 12,831 lncRNA genes (Figure 5E). Remarkably, if we reconsider the number of isoforms from Figure 1B that have unique or partially unique junctions that can be targeted with a predicted active junction-spanning gRNA, we conclude that up to 89% of isoforms from multi-isoform genes (excluding the 5,195 protein-coding and 11,187 lncRNA single isoform genes) are uniquely targetable (Figure 5F). These targetable isoforms harbor all types of splicing and transcriptional events, including cassette exons, alternative 5’ and 3’ SS, and alternative starts and ends. These findings solidify our strategy to target gRNAs to exon-exon junctions as a versatile method that can be broadly applied to knockdown specific isoforms and study their function.

## Discussion

Driven in part by the widespread adoption of RNA sequencing and accompanying algorithms for differential splicing and splicing QTL mapping, it is increasingly clear that AS (and isoform variation more broadly) is a major player in development, cell-type specific regulatory programs, evolution, and disease, including Mendelian conditions, cancer and complex traits such as neuropsychiatric disorders. However, it is challenging to study the function of implicated splice isoforms in an endogenous cellular context with existing experimental and computational methods. In this paper, we make significant progress to address this challenge.

We introduce a strategy for isoform-specific RNA knockdown that targets Cas13/gRNAs to exon-exon junctions in the mature RNA product. A similar approach has been used previously to target back-spliced junctions of circRNAs with Cas13 (Zhang et al. 2021) but has not been extensively explored for normal splice junctions. Our method offers several distinct advantages to other approaches to perturb isoform expression. We show that our strategy can specifically knockdown isoforms corresponding to all major event types, including alternative first and last exons, alternative 5’ and 3’ SS, and both inclusion and exclusion products from a cassette exon, while other methods can only target one type of event (e.g. Cas9, Cas9-Cas12a, Cas13, RNAi, or ASOs targeting a cassette exon inclusion). In addition, targeting junctions ensures that the pre-mRNA is not knocked down, which would decrease overall gene expression. We recognize the potential for feedback mechanisms to affect gene and isoform expression upon RNA knockdown with Cas13, but this would be true for any RNA-targeting approach. However, at least for the examples in this paper we do not see strong compensatory changes. In particular, for *RBFOX2*, which is well-known to bind and regulate its own transcripts (Damianov and Black 2010), we observe a modest increase in the TPM of the *untargetted* isoforms. In summary, our strategy provides the most versatile platform currently available for isoform-specific knockdown.

Previous Cas13 gRNA efficacy prediction models trained using exon-targeting gRNAs have found that gRNA efficiency is primarily dependent on gRNA and target RNA sequence. We find that this holds true for junction gRNAs, and that the sequence determinants are indistinguishable to those for the previously investigated exon gRNAs. We did not detect any effect of junction sequence on gRNA efficacy beyond the gRNA/target sequence itself, possibly implying that RBP binding does not play a substantial role in determining gRNA efficacy. We did briefly explore explicitly including predicted or measured (in other cell-types) RBP binding as additional features, but saw no improvement in predictive accuracy (results not shown). We caution against over-interpreting this finding however given the limited cross cell-type accuracy of existing RBP binding predictors, and the lack of RBP binding profiles in A375. Taken together, we speculate that gRNA design for junctions and exons will be similar across other RNA targeting CRISPR/Cas systems and across other species (Kushawah et al. 2020).

When designing an isoform-centric experiment, an initial challenge is confidently identifying which isoforms are expressed in the cell type of interest. Short read RNA-sequencing is problematic due to the challenges of ambiguous read mapping and transcript assembly. In our experience, LRS results in more accurate measurements than any obtained from short read RNA-seq to determine the presence or absence of specific isoforms. In our PacBio data, we detect a significant number of unannotated isoforms, consistent with other studies (Reese et al. 2023). For example, we were surprised to find that the predominant *RBFOX2* isoform containing the most upstream alternative start junction (RBFOX2.178149) was an unannotated isoform and not *RBFOX2-220*, the canonical isoform, or *RBFOX2-209*. Especially when interpreting isoform knockdown results, it is critical to know with confidence which isoform(s) are being perturbed. Importantly, Isoviz can incorporate both long and short read data, using long reads to identify isoform structures and short reads to give additional quantitative support at the junction-level.

In this paper, we do not measure protein-level changes upon RNA knockdown. Identifying and quantifying protein isoforms using Mass Spectrometry (MS) has similar limitations to identifying RNA isoforms with short read sequencing data alongside additional technical challenges. It is difficult to assign fragmented peptides to the correct isoform, and some isoforms may not have a unique peptide that can be detected with MS (Miller et al. 2022; Reixachs-Solé and Eyras 2022). This remains an important area for future work to link RNA and protein isoform expression and function.

TIGER (Wessels et al. 2023) gRNA efficacy predictions are highly correlated with RNA knockdown. While our essentiality screen had many inactive guides due to its intentionally unbiased design, Isoviz combined with TIGER will enable the design of efficient junction-targeting gRNAs for both single gene experiments and high-throughput screens to interrogate isoform-specific functions for phenotypes of interest.

## Methods

### Cas13d Essentiality Screen

#### Monoclonal cell line generation for screen

A375 cells were acquired from American Type Culture Collection (ATCC Cat #CRL-1619). A375 cells were maintained at 37°C with 5% CO2 in DMEM media (ThermoFisher Cat #11965118) supplemented with 10% serum (Serum Plus II, Sigma-Aldrich, Cat #14009C).

To generate the doxycycline-inducible Cas13d A375 cell line used in the screen, we transduced cells with lentivirus carrying the previously published plasmid, pLentiRNACRISPR_007 (Addgene #138149) at a low multiplicity of infection (MOI <1). Cells underwent drug selection for one week with 5ug/ml of blasticidin (ThermoFisher, Cat #A1113903). Colonies were picked after sparse plating, expanded, dox induced, and screened for Cas13d expression using western blotting and immunofluorescence methods with an anti-HA antibody (Cell Signaling Cat #2367). Cas13 knockdown activity was confirmed by FACS using CD46 as a positive control (Cell Signaling Cat #13241).

#### Guide library design

To design the Cas13d gRNA library targeting exon-exon junctions in A375 melanoma cells, we identified junctions that have evidence of expression. First, we downloaded three publicly available RNA-seq datasets from A375 cells (SRR6515912, SRR6515913, and SRR6515914 from GSE109731; (Regan-Fendt et al. 2019). Next, we processed sequencing reads according to LeafCutter recommendations (Y. I. Li et al. 2017): reads were aligned to hg38 using STAR v2.7.1 with parameters ‘--twopassMode’ and ‘--outSAMstrandField intronMotif’ (Dobin et al. 2013). Sequencing reads that overlap exon-exon junctions were counted using ‘regtools junction extract’ v0.5.2 with parameters ‘-a 8 -m 50 -M 500000 -s 1’ (Cotto et al. 2023). Next, to define intron clusters using LeafCutter, we ran the ‘leafcutter_cluster_regtools.py’ python script with ‘-m 30 -p 0.01 -l 500000 parameters’ (Y. I. Li et al. 2017).

To select genes to target in our essentiality screen, we employed a multi-step filtering approach. First, we considered 1000 candidate genes that had the lowest LFC (i.e. genes that, when knocked out, reduced cell proliferation) from the GeCKO essentiality screen using CRISPR/Cas9 in A375 cells (Shalem et al. 2014). Using the GENCODE v41 basic annotations as a reference, globally, we defined junctions present in all transcripts of a gene (common), present in a subset of transcripts (partially unique), or present in only one transcript of a gene (fully unique). From our 1000 initial candidate essential genes, we selected two common junctions with the highest read counts per gene and partially or fully unique junctions that LeafCutter clustered into groups of twos (likely alternative 5’ ss or 3’ss), threes (likely cassette exons), or fours. We also required unique junctions to have at least 15% usage across all three replicates. Next, we designed eight 23bp gRNAs overlapping each junction, starting at -15bp and +8bp and sliding by 1bp to -8bp and +15bp relative to the splice junction. We chose this tiling window of up to 15bp overlapping a single exon based on previous work showing loss of RNA knockdown efficiency using gRNAs less than 20bp (Wessels et al. 2020). We chose this conservative 15bp cutoff, but others could test 16/7, 17/6, 18/5, and 19/4 splits to see if they can achieve isoform-specific RNA knockdown. We used ‘bedtools fasta’ to get the sequence.

To obtain the final set of gRNAs, we used Bowtie (Langmead et al. 2009) to filter out gRNAs that aligned to more than one gene locus when allowing up to 2 mismatches. Additionally, we eliminated gRNAs with homopolymers with a length 5 or more for A’s, G’s, and C’s and 4 or more for T’s. After filtering, we had 50,512 gRNAs targeting 6,957 junctions in 837 GeCKO-defined essential genes. Additionally, as controls, we included 963 non-targeting guides (random 23bp sequences with no matches in the transcriptome) and 1,992 common junction gRNAs targeting 413 non-essential genes.

#### gRNA library synthesis, cloning, and amplification

Pooled gRNA libraries were synthesized as single-stranded oligonucleotides (Twist Biosciences) and resuspended to a concentration of 10ng/µl in TE. Cloning and sequencing were performed the same as by Wessels and colleagues (Wessels et al. 2023).

Briefly, overhangs for Gibson cloning were added to the oligonucleotide library by PCR using the oligo_amp_FW and oligo_amp_REV primers (Supplementary Table S1). Libraries were PCR amplified in 8x 50ul reactions per 10,000 gRNAs (0.5 µl Q5 polymerase (NEB Cat #M0493), 10 µl 5× reaction buffer, 2 µl oligo pool (1 ng/µl), 2.5 µl of each forward and reverse primer (10 µM), 2.5 µl dNTPs (10 mM) and 30 µl water). PCR conditions were 98 °C/30 s, 10× (98 °C/10 s, 63° C/10 s and 72 °C/15 s) and 72 °C/3 min. The PCR amplified library was gel-purified, quantified, and cloned into BsmBI-digested pLentiRfxGuide-Puro (Addgene #138151) via Gibson Assembly. Eight Gibson reactions were performed with a 20-µl reaction volume each time (500 ng digested plasmid (0.088 pmol), 123.15 ng purified oligo pool (1.3245 pmol, 15:1 molar ratio), 10 µl 2× Gibson Assembly Master Mix (NEB)), incubated for 1 h at 50 °C. Next, to expand the gRNA library, the assembled library plasmid pool was electroporated into Endura cells (Lucigen, Cat #60242-2) at 50-100 ng/µl. After electroporation, cells were recovered in LB medium for 1 hour and plated on LB agar carbenicillin at 37°C for 12-14h. To achieve good library representation, we aimed to get a coverage of > 200 colonies per gRNA. The library plasmid pool was extracted from harvested bacterial cells with the IBI Maxiprep Kit (Cat #IB47125). Complete library representation with minimal bias was verified by Illumina sequencing (MiSeq, Cat #MS-103-1002).

#### gRNA library screening and sequencing

Lentivirus was produced via transfection of library plasmid pool and appropriate packaging plasmids (psPAX2, Addgene #12260 and pMD2.G, Addgene #12259) using linear polyethylenimine MW25000 (Polysciences, Cat #23966). We seeded ten million A375 cells per 10 cm dish and transfected with 60 µl polyethylenimine, 9.2 µg plasmid pool, 6.4 µg psPAX2 and 4.4 µg pMD2.G. At 3 d post-transfection, viral supernatant was collected and passed through a 0.45-µm filter and stored at −80 °C until further use.

Doxycycline-inducible RfxCas13d-NLS A375 cells were transduced with the pooled library lentivirus in separate two infection replicates, ensuring at least 1,000× guide representation in the selected cell pool per infection replicate using spinfection. After 24 h, cells were selected with 1 µg ml−1 puromycin (ThermoFisher, Cat #A1113803), resulting in ∼30% cell survival. Puromycin selection was performed 72 h after the addition of puromycin. Assuming independent infection events (Poisson), we determined that ∼83% of surviving cells received a single sgRNA construct (Sidi Chen et al. 2015). After completed puromycin selection, input sample was collected (Day 0), and RfxCas13d expression was induced by the addition of 1 µg/ml doxycycline (Sigma-Aldrich, Cat #D9891). Cells were passed every 2–3 d (maintaining full representation) and supplemented with fresh doxycycline. We collected genomic DNA (gDNA; at least 1,000 cells per construct representation) from each sample on Day 0, Day 7, Day 14, and Day 21.

To extract gDNA, screen cells were lysed as in (Sidi Chen et al. 2015) with 12 ml of NK lysis buffer for 100 million cells (50 mM Tris, 50 mM ethylenediaminetetraacetic acid, 1% SDS and pH 8). Once cells were resuspended, 60 µl of 20 mg/ml Proteinase K (Qiagen) was added and the sample was incubated at 55 °C overnight. The next day, 60 µl of 20 mg/ml RNase A (Qiagen) was added and mixed, and samples were incubated at 37 °C for 30 min. Then, 4 ml of prechilled 7.5 M ammonium acetate was added, and samples were vortexed and spun at 4,000g for 10 min. The supernatant was placed in a new tube, mixed well with 12 ml isopropanol and spun at 4,000g for 10 min. DNA pellets were washed with 12 ml of 70% ethanol, spun and dried, and pellets were resuspended with 0.2× TE buffer (Sigma-Aldrich). In addition, we also generated linearized plasmid library input and diluted it down to mimic similar copy number conditions as the gDNA samples.

To amplify the gRNA cassette from gDNA and add indexing for Illumina sequencing, we used a two-step PCR protocol, PCR1 and PCR2, respectively. For the PCR1 reaction, we used 960 µg gDNA for each sample. We performed 96× 100 µl PCR1 reactions per sample (10 µl 10× Taq buffer, 0.02 U/µl Taq-B enzyme (Enzymatics, P7250L), 0.2 mM dNTPs, 0.2 µM forward and reverse primers and 100 ng gDNA per µl). Thermocycler conditions were 94 °C/30 s, 20× (94 °C/10 s, 55 °C/30 s, 68 °C/45 s) and 68 °C/3 min. For each sample, all PCR1 products were pooled and mixed. For the PCR2 reaction, we performed 2 reactions per sample (20 µl 5× NEB Q5 buffer, 0.01 U/µl Q5 enzyme, 20 µl PCR1 product, 0.2 mM dNTPs and 0.4 µM forward and reverse PCR2 primers in 100 µl). Thermocycling conditions for PCR2 were 98 °C/30 s, 7× (98 °C/10 s, 63 °C/30 s and 72 °C/45 s) and 72 °C/5 min. For each sample, PCR2 products were pooled, followed by normalization (gel-based band densitometry quantification), before combining equal amounts of uniquely barcoded samples.

The pooled product was then purified using SPRI beads. First, we performed a 0.6× vol/vol SPRI to remove gDNA carryover, followed by the addition of a 0.3× vol/vol SPRI (0.6 + 0.3 = 0.9× final) to the supernatant to purify the ∼260 bp PCR product. The final amplicons were sequenced on Illumina NextSeq 500-II HighOutput 1 × 150 v2.5 (Cat #20024907).

Oligonucleotides can be found in Supplementary Table S1.

### Cas13d Essentiality Screen Analysis

#### Data Pre-processing

Cutadapt v3.5 (Martin 2011) was used to demultiplex reads based on barcode sequences in the forward primer of PCR2 during screen library prep (Supplemental Table S3). We included the 8bp barcode plus 8bp of the U6 sequence (“TCTTGTGG”) to ensure the proper position of the barcode match within the read. We allowed for 1 mismatch within the 16bp sequence (-e 1 -O 16, – action=none). Cutadapt was also used to trim 5’ (54bp) and 3’ (16bp) sequences upstream and downstream of the gRNA sequence (-e 0.1). Bowtie v1.1.2 (Langmead et al. 2009) index was built from the screen library fasta file and reads were subsequently aligned to the library index with strict parameters of -v 0 that do not allow for mismatches and -m 1 that only allow for a single alignment (--norc --best --strata). We used an in house R script that inputs the sam alignment file and outputs a raw count matrix of gRNA read counts per sample.

Technical replicates, re-sequencing of the same library for greater read depth, were combined for each sample and all future analyses were performed on this pool. Then, we added 1 to all values to prevent division by 0 for entirely depleted gRNAs. We divided counts for each guide by its geometric mean (across conditions) before normalizing by the median of ratios (Love, Huber, and Anders 2014) corresponding to the non-targeting gRNAs. We visualized normalized counts and removed gRNAs that had low or high detection in the Day 0 sample (cutoffs are 0.01 and 0.99 quartiles of normalized gRNA count distribution). Finally, we calculated log2 fold changes (LFCs) by dividing normalized counts across each time point by the corresponding replicate at Day 0 and taking their log2.

#### Re-defining Cas13d essential genes

To label gRNAs as active vs inactive, we employ the same methodology as (Wessels et al. 2023). We fit a normal distribution to the mean of the non-targeting gRNAs’ replicates using maximum likelihood estimation. We then define an active gRNA whose LFC falls below the 1% of this distribution.

After labeling all gRNAs in our screen as active or inactive, we compute the per-gene ratio of active gRNAs. We require that, to be defined as essential, a gene must have at least 5% of its gRNAs classified as active.

#### Comparison to previous essentiality screens

We compared gene essentiality scores from our screen to scores from four previously published essentiality screens in A375: GeCKO (CRISPRko), Brunello (CRISPRko), Dolcetto (CRISPRi), and RNAi. For GeCKO, we obtained an initial dataset from the Cancer Dependency Map (DepMap, file: achilles_geckoV2_19Q4.csv.gz). This dataset contained a single LFC value for every gene. For Brunello, we obtained a raw count matrix from the published A375 genome-wide negative selection screen (Doench et al. 2016). LFCs were averaged across both replicates and then for all gRNAs corresponding to the same gene to get a single score per gene. From the Dolcetto A375 screen, we similarly obtained a matrix of raw counts (Sanson et al. 2018). This screen contained three replicates and LFCs were averaged across replicates and gRNAs per gene. For the RNAi screen in A375, data was also downloaded from DepMap (file: D2_combined_gene_dep_scores.csv). A single score indicating LFC was provided for every gene.

For the purpose of comparing gene essentiality scores across all screens, we only included genes assessed in all four publicly available screens as well as in our Cas13d screen. This resulted in 1,165 genes. To calculate gene essentiality scores from the Cas13 screen date, we take the median LFC of the top four gRNAs per junction and then the median LFC across the top four junctions to get a single value per gene. We used the ggcorrplot R package to assess the Pearson correlation across the A375 essentiality screens. We evaluated the correlations with individual time points and replicates in our Cas13 screen.

#### SEABASS Linear Mixed Model

Existing approaches for analyzing essentiality screen data do not handle multiple time-points and replicates. We developed a probabilistic, hierarchical linear mixed modelling approach which we call SEABASS (Screen Efficacy Analysis with BAyesian StatisticS). We model log2 fold changes *y_gtr_* for guide *g* at time-point (week) *t* and replicate *r* using the linear mixed model,

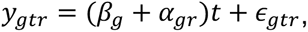

where *β_g_* is a per guide slope, *∈_gtr_* ∼ *T*(0, *σ*_1_, *τ*_1_) is noise, *⍺_gr_* ∼ *T*(0, *σ*_2_, *τ*_2_) are random slope (i.e. random effect) terms, and *T*(*m*, *σ*, *τ*) is a Student-t distribution with mean *m*, scale *σ*, degrees of freedom *τ* and therefore variance *τσ*^2^/(*τ* − 2) (for *τ* > 2). The hyperparameters {*σ*_1_, *τ*_1_, *σ*_2_, *τ*_2_} are shared across all guides (and genes). By fitting *τ*_1_ and *τ*_2_ we can control how heavy-tailed the noise distribution is. For *τ* = 1 the Student-t corresponds to a Cauchy distribution (extremely heavy tails), and for *τ* → ∞ a Gaussian (light tails). The noise distribution parameters {*σ*_1_, *τ*_1_, *σ*_2_, *τ*_2_} are learnt on the non-targeting (NT) guides only, where we fix *β_g_* = 0. We additionally explored using Laplace or explicit Gaussian or Cauchy distributions for the noise and random slope terms but found these provided a worse fit to the NT data according to the evidence lower bound (ELBO). Our use of the Student-t distribution endows SEABASS with natural robustness to outliers.

We put a gene-dependent prior on the per-guide slopes *β_g_* ∼ *D*(0, *s*_*γ*_) where *D*(*m*, *s*) is a location-scale distribution with mean 0 and scale *s*, and *γ* is the gene targetted by guide *g*. The per gene scales *s*_*γ*_ capture differences in gene essentiality. We explored *D* being Gaussian, Cauchy, Student-t or Laplace, and choose Laplace since it gave the lowest estimated false positive rate (assuming all significantly positive *β_g_* are false positives). We put a log-normal prior on *s*_*γ*_ i.e. 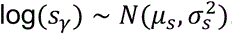.

We use stochastic variational inference (SVI) in pyro (https://pyro.ai/) to fit the model jointly across all guides (and all genes). We use a structured variational posterior where {*β_g_*, *⍺_g_*_1_, ⋯, *⍺_gR_*} for a guide are drawn from a multivariate normal (where *R* is the number of replicates). This is to account for the strong posterior dependencies we expect between these variables. For *s*_*γ*_ we use a (diagonal/mean-field) normal variational posterior on each log(*s*_*γ*_). We optimize {*σ*_1_, *τ*_1_, *σ*_2_, *τ*_2_, *μ*_2_, *σ*_’_} by placing Dirac delta variational “posteriors” on these parameters.

We found that SVI using just one Monte Carlo (MC) sample for gradient estimation (the default in pyro) and the Adam optimizer did not fully converge. To address this, we developed a strategy where we monitored the ELBO for optimization having stalled, assessed by the ELBO for the last 10 epochs not showing a statistically significant (p>0.05) downward slope. We then double the number of MC samples used for gradient estimation and resume optimization. We go up to a maximum of 32 MC samples. This resulted in improved ELBOs and agreement between parameter estimates across random initializations.

#### TIGER Model Architecture

We closely mimicked the published TIGER architecture (Wessels et al. 2023): two convolutional layers with 32 filters, each of length 4, and rectified linear unit (ReLU) activations. This is followed by a max pooling layer with a pool size of 2. The data is then flattened into a vector and subjected to dropout with a rate of 0.25 for regularization. Subsequently, the features are concatenated with non-sequential features and fed into a dense layer with 128 sigmoid outputs. Another dropout layer with a rate of 0.1 is applied here. This is succeeded by a dense layer with 32 sigmoid outputs and another dropout layer with the same 0.1 rate. The final layer is a linear output layer, producing a scalar LFC prediction. We additionally score these predictions (i.e. map them to the unit interval) using a sigmoid function similar to (Wessels et al. 2023; Wei et al. 2021; Cheng et al. 2023). In particular, we map the 0.1 quantile (very active LFCs) of all gencode LFC predictions to 0.9 (a high score) and the 0.9 quantile (inactive LFCs) of all gencode LFC predictions to 0.1 (a low score).

In generating TIGER’s, TIGERjunctions’s, and TIGERbass’s predictions for our screen, all gencode junctions, and our RT-qPCR tested gRNAs, we performed 10-fold cross validation, ensuring that any gencode or RT-qPCR gRNA never appears in the training pool. This procedure results in a single prediction for gRNAs that appear in the training set TIGER uses (Wessels et al. 2023). TIGERjunction uses our screen data, and TIGERbass uses SEABASS slope estimates from our screen data–and an ensemble of 10 predictions for gRNAs that did not, which we then average. TIGERjunction and TIGERbass only consider transcript sequence since guide sequence is redundant in our setting of perfectly matched gRNAs. For TIGER guide models (Figure 2, Supplemental Figure 2), we use +/-1 nucleotide of additional transcript sequence context like the published TIGER architecture (Wessels et al. 2023). However, we do not normalize for gene essentiality as TIGER does, since this requires a significant number of guides targeting a single gene, which our screen does not have.

In the junction-targeting setting, Wessels et al. (2023)’s additional scalar features related to gRNA position along a transcript, distance to nearby junctions, and transcript length are ill-defined. Their results suggest these features have minimal impact on predictions (via SHAP analysis) and on predictive performances (via feature holdouts). Therefore, we only consider the following features for our training of TIGER, TIGERjunction, and TIGERbass (Figure 2, Supplemental Figure S2):

- Target accessibility: log unpaired probability, log unpaired probability at positions 11, 19 and 25
- Hybridization MFE at positions 1 and 23, 2 and 12 and 15 and 9
- Guide MFE
- Guide secondary structure

○ presence of direct repeat stem loop (a Boolean variable)
○ presence of a g-quadruplex (a Boolean variable)

For TIGERsite (Figure 2), we provide 100 nt of transcript sequence only and do not consider these additional scalar features.

#### Assessing Model Feature Importance

To determine TIGER’s learned gRNA design rules, we performed ten-fold cross validation, collecting Shapley additive explanations (SHAP) (Lundberg and Lee 2017) values for each element in the fold such that we have a SHAP value for every dataset element. Averaging these values conditioned on positional nucleotide identity, we observed TIGER learns known gRNA design rules both when trained on exon data (Wessels et al. 2023) and when trained on our junction screen data. We similarly collected SHAP values for the junction-sequence TIGER model (Figure 2D) that predicts the average of a junction’s tiled gRNAs’ LFCs from 50 nt up- and down-stream the splice junction.

#### Comparison to other publicly available models

To obtain Cheng et al. predictions for all junction-spanning gRNAs, we re-trained the DeepCas13 model using LFCs for the 5,726 gRNAs provided in their paper (Cheng et al. 2023)(https://bitbucket.org/weililab/deepcas13/src/master/). We then used this model to predict gRNA efficiency for all 2.2 million GENCODE junction-spanning gRNAs. DeepCas13 outputs slightly different but highly correlated (r > 0.94) predictions each time it is trained. Therefore, we repeated this process 5 times and averaged them to get a single prediction score per gRNA. To obtain Wei et al. predictions for all junction-spanning gRNAs, we downloaded the already generated predictions for Human RefSeq coding genes [refseq_coding_guides_prediction_sorted.csv] from https://www.rnatargeting.org/ (Wei et al. 2021) and merged the 7.4 million gRNA predictions with our GENCODE junction gRNA list. Their model uses gRNAs that are 30bp long, so we extracted the first 23bp. This results in gRNA predictions for 1,617,364 of the 2.2 million GENCODE gRNAs.

#### Analysis of non-sequence feature association with gRNA efficacy residuals

We considered certain non-sequence features for gRNAs and junctions to evaluate whether they may help explain discrepancies in gRNA efficacy predictions beyond what is captured by TIGER_bass_. Guide-level residuals were calculated as observed slopes minus predicted slopes, and junction-level residuals were calculated by averaging all available gRNAs per junction. The number of guides, junctions and genes evaluated herein included 27,804, 3,822 and 480 respectively.

For non-sequence features, some datasets were generated in-house, while others were pulled from additional sources.

A375 gene expression, percent gene nuclear, and percent junction nuclear were calculated from A375 RNA-seq data generated in this paper (see ‘Methods: Illumina short read sequencing analysis’ and Supplemental Table S3 for additional information). Briefly, gene counts were obtained using featureCounts (Liao, Smyth, and Shi 2013) and were normalized to RPKM (Reads Per Kilobase Million) and junction counts were obtained using RegTools (Cotto et al. 2023). Percent nuclear values were obtained by dividing the nuclear read counts by the sum of the nuclear and cytoplasmic read counts for a gene or junction. We recognize that these are not absolutely quantifications of nuclear localization but are relative quantifications for comparison between genes or junctions.

Gene length and intron length were obtained from Hg38 coordinates from the ‘annotables’ package (https://github.com/stephenturner/annotables). Intron lengths were calculated using observed junction lengths. gRNA tiling position and relative junction position were obtained from GENCODE v41 annotations. gRNA tiling position indicates positions 1-8 along a junction where 1 is -15bp/+8bp and 8 is -8bp/+15bp. Relative junction position was calculated based on a junctions’ basepair genomic distance to the gene TSS.

To obtain mean RNA half lives, we downloaded a matrix of genes and RNA half lives across 33 studies (Agarwal and Kelley 2022). Half lives were averaged for all available studies for each gene.

All available features were centered and scaled to ensure that all predictor variables were on a similar scale. We then fit a multiple linear regression to assess each feature’s regression coefficient, standard error and p-value.

### piggyBac-Cas13 Knockdown Experiments

#### Cloning of Cas13d and gRNA constructs into piggyBac transposon system

To create the doxycycline-inducible Cas13d piggyBac vector, the TRE_NLS-*Rfx*Cas13d-NLS-HA was cloned from Addgene plasmid # 138149 (Wessels et al. 2020) into the piggyBac backbone from Addgene plasmid # 126029 (Schertzer et al. 2019) by digestion with SpeI and EcoRI followed by Gibson Assembly (NEB).

To create the rtTA-sgRNA expressing piggyBac vector, the hU6_*Rfx*Cas13d-DR_BsmBI was cloned from Addgene plasmid # 138151 (Wessels et al. 2020) in the piggyBac backbone from Addgene plasmid # 126028 (Schertzer et al. 2019) by digestion with SfiI and BglII followed by Gibson Assembly (NEB). Oligonucleotides used for cloning are in Supplemental Table S1. Full plasmid sequences were verified using plasmidsaurus long read sequencing (https://www.plasmidsaurus.com/).

We plan to submit both plasmids to Addgene after publication: (1) PB_TRE_NLS-*Rfx*Cas13d-NLS-HA and (2) PB_hU6_*Rfx*Cas13d-DR_BsmBI.

To allow propagation of the piggyBAC transposase from System Biosciences on ampicillin plates, the transposase was cloned into SmaI and HindIII sites into pUC19 (NEB) as in (Kirk et al. 2018).

#### Cell culture

HEK293 cells were acquired from ATCC (Cat #CRL-1573). HEK293 cells were maintained at 37°C with 5% CO2 in DMEM media (ThermoFisher Cat #11965118) supplemented with 10% serum (Serum Plus II, Sigma-Aldrich, Cat #14009C).

HUES66 human embryonic stem cells (hESCs) were obtained from Harvard University. Cells were maintained at 37°C with 5% CO2 in StemFlex media (ThermoFisher Cat #A3349401) and grown on Geltrex-coated plates (ThermoFisher Cat #A1413302). For passaging, Accutase (ThermoFisher Cat #A1110501) was used to dissociate cells and 10 µM ROCK inhibitor Y-27632 dihydrochloride (Tocris Cat #1254) was added to the media for plating.

#### Stable transfections of piggyBac Cas13 system

With piggyBac, the number of integrations of cargo plasmids can be tightly controlled by changing the ratio of cargo plasmids to PB transposase plasmid (Cadiñanos and Bradley 2007; Wilson, Coates, and George 2007; Schertzer et al. 2019). We initially tested two gRNAs at five different gRNA:Cas13:transposase ratios in human embryonic stem cells (hESCs), but we did not observe significant differences in RNA knockdown (data not shown). To minimize the number of insertions, we perform all experiments at a 2:2:1 ratio.

To generate stable Cas13/gRNA-expressing HEK293 cell lines, 2 x 10^5^ cells were plated in a single well of a 12-well plate and transfected the following day using Lipofectamine 3000 (Invitrogen #L3000001) according to the manufacturer’s protocol. A 2:2:1 ratio of piggyBac cargo vectors (Cas13d and gRNA) and pUC19-piggyBac transposase, totaling 1.25 µg total of plasmid DNA was mixed with Mix #1 (50 µl OptiMEM + 2.5 µl P3000) per well. Mix #2 (50 µl OptiMEM + 1.88 µl Lipofectamine 3000 reagent) was added and incubated for 10 minutes before adding to individual wells. On the day after transfection, cells were selected with 100 µg/mL Hygromycin (ThermoFisher #10687010) and 400 µg/mL G418 (ThermoFisher #10131035) for 7 to 9 days. After selection, Cas13d expression was induced by adding 1 µg/mL doxycycline (Sigma #D3447-500MG) to the media for 24 hours before cells were harvested.

#### RNA Extraction and RT-qPCR analysis

RNA was extracted using TRIzol (Invitrogen #15596018) according to the manufacturer’s protocol. The Qubit RNA HS Assay Kit (Invitrogen #Q32855) was used to quantify RNA and the High Capacity cDNA Reverse Transcription Kit (Invitrogen #4368813) was used to RT 500 ng - 1 µg of RNA. qPCR was performed using iTaq Universal SYBR Green (Bio-Rad #1725122) and primers listed in Supplemental Table S1.

#### RNA Fractionation

To isolate RNA from cytosolic and nuclear fractions, cells were washed twice with 1mL cold PBS and scraped in 1mL PBS + 1mM PMSF + 1:100 protease inhibitor cocktail (PIC, Sigma P8340). 200uL was removed at this step and added to 1mL Trizol (total RNA). The remaining cells were centrifuged at 1500xrcf. for 5 min, and resuspended in 250uL low salt solution (10mM KCl, 1.5mM MgCl2, 20mM Tris-HCl pH 7.5) supplemented with 1mM PMSF, 1mM DTT, and 1x PIC. Triton X-100 was added to a final concentration of 0.1% and cells were rotated for 10 min at 4C, then centrifuged for 5 min at 1500xrcf. 200uL of supernatant was removed and added to 1mL Trizol (cytosolic fraction). The remaining supernatant was discarded and the nuclear pellet was washed by rotating for 2min at 4C in low salt solution without Triton X-100 and centrifuged at 1300rpm for 10 min. Nuclear pellet was resuspended in 1mL Trizol (nuclear fraction). Isolation of RNA from Trizol was performed according to the manufacturer protocol.

### High Throughput Sequencing and Additional Analyses

#### Analysis of GENCODE basic annotations

The R code used to generate the GENCODE summary barplots in Figure 1B (human) and Supplementary Figure 1A are added to the Isoviz package (isoviz_junction_to_transcript_summary.R). The code can be run in two ways: 1) using GENCODE annotations or 2) long read sequencing generated annotations specific to a cell type. GENCODE annotations for additional species beyond human and mouse may also be used. The table includes all junctions present in the input file with two classification columns: ‘junction_category’ indicates whether the junction is classified as common, fully unique, partially unique, or single isoform and ‘Isoform_targetable’ indicates whether the isoform can be targeted uniquely. When comparing classifications of junctions and isoforms using long read data annotations, outputs are expected to vary based on cell type. It is recommended to consider junction and transcript classifications specific to your cell type of interest when designing experiments.

#### Illumina short read sequencing

Per RNA sample, we combined 1 µg of total RNA + 2 µl 1:100 ERCC RNA Spike-in Mix #1 (Invitrogen #4456740). Libraries were prepared using the Kapa RNA HyperPrep Kit with RiboErase (Roche #08098131702), pooled, and sequenced using the Nextseq 500/550 High Output Kit with 150 cycles (Illumina #20024907).

#### Illumina short read sequencing analysis

RNA-seq reads were aligned to hg38 genome using STAR v2.7.1 and ‘--twopassMode Basic’ and ‘--outSAMstrandField intronMotif’ parameters (Index built with GENCODE v41 basic annotations gtf (Dobin et al. 2013)). Samtools v1.9 was used to filter for reads with a mapping quality greater than 20, sorted, and indexed (H. Li et al. 2009).

To determine the gene-level RPKM and TPM counts in A375 total, nuclear, and cytoplasmic RNA-seq data, alignment bam files were overlaid with the GENCODE v41 basic annotation gtf file using featureCounts (subread v2.0.4) (Liao, Smyth, and Shi 2013).

HEK293, hESC, and A375 junction counts (provided in Isoviz and used for non-sequence features in Supplementary Figure S2) were generated using Regtools extract v0.5.2 (Cotto et al. 2023) with ‘-a 8 -m 50 -M 500000 -s 0’ parameters.

#### PacBio long read sequencing

Prior to sending RNA samples to PacBio for long read sequencing, the RNA integrity number (RIN) was calculated using the RNA 6000 Nano Kit (Agilent #5067-1511). RIN scores ranged between 8.7-9.7, indicating high quality, intact RNA. Per RNA sample, we combined 1 µg of total RNA + 2 µl 1:100 ERCC RNA Spike-in Mix #1 (Invitrogen #4456740).

MAS-Seq bulk Iso-Seq libraries were generated in collaboration with PacBio using a pre-commercial protocol that implements the MAS-Seq concatenation method (Al’Khafaji et al. 2023). Briefly, cDNA molecules were concatenated into an ordered array, in this case, an 8-fold array, and sequenced on the Revio system. The concatenated array is sequenced as a HiFi read and then bioinformatically de-concatenated into segmented reads (S-reads) which represent the original cDNA molecules.

#### PacBio long read sequencing analysis

A pre-commercial version of the SMRT LINK analysis pipeline was used to generate HiFi reads, S-reads, and subsequent full-length reads and high-quality isoform sequences. After the *isoseq3 refine* step where full-length non-chimeric reads were obtained in bam file format, we continue with our own analysis pipeline.

Minimap2 v.2.17 (H. Li 2018) was used to align reads to GRCh38 human genome reference with -ax splice:hq -uf parameters. Secondary alignments were removed, and aligned reads were filtered for mapping quality greater than or equal to 60. The resulting bam files were converted to BED12 format and PSL format with FLAIR’s helper scripts (Tang et al. 2020); https://flair.readthedocs.io/en/latest/cite.html). Then, additional filtering was applied to select for full length reads based on overlap with annotated transcription start sites (TSS) and transcription end sites (TES). Our custom script required overlap between -10bp to +50bp of TSS and -50bp to +10bp of TES (using human GENCODE v41 Basic annotation). Reads that shared the same junction chain were grouped together and collapsed, using a custom script adapted from FLAIR collapse (Tang et al. 2020). Importantly, our custom collapse script ignores start and end coordinates of the reads, allowing reads with the same junction chain, but slight variations in TSS or TES, to be grouped together. Unspliced reads (i.e. mono-exonic reads) were excluded. Resulting full length, collapsed reads were matched with transcripts in GENCODE v41 basic annotations, and any transcripts without a match were still overlapped with the proper gene and was assigned a unique transcript ID through our custom scripts. Finally, the number of reads grouped together per transcript were tallied and output to a separate file per sample. The “million” in TPM was calculated based on the total number of full length reads after filtering.

## Supplemental Figures

**Supplemental Figure 1.**
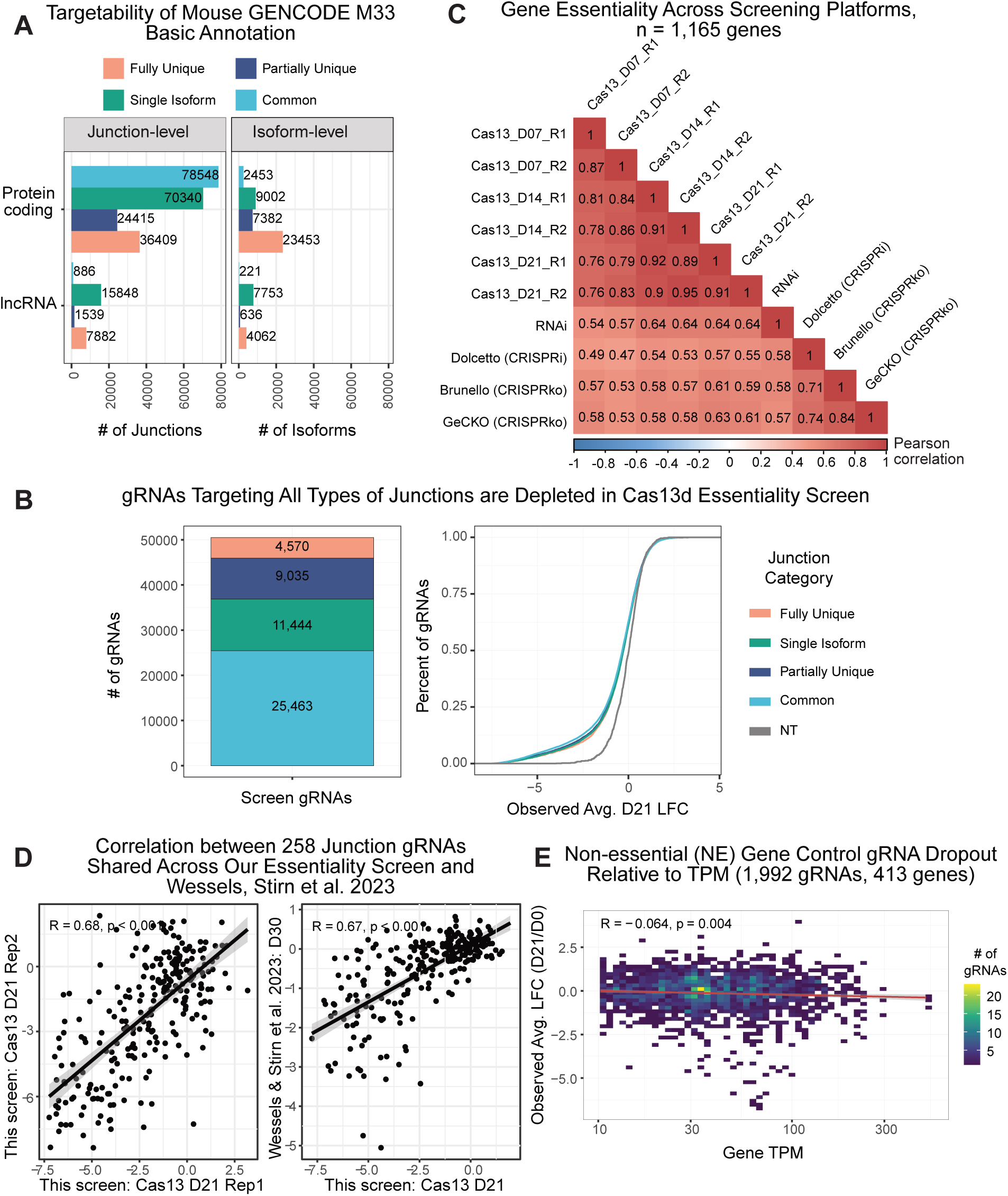
(A) Counts of junction-level and isoform-level categories described in Figure 1A for the GENCODE mouse vM33 Basic Annotation (analogous to Figure 1B). (B) Barplot (left) shows the number of gRNAs in our essentiality screen that target common and single isoform junctions (targets all isoforms a gene) and fully unique and partially unique junctions (targets a subset of isoforms). Cumulative density function plot (right) comparing Day 21 LFC of gRNAs targeting different junction types. Number of gRNAs per category is given in the barplot on the left. NT is used as a reference (gray line). (C) Pearson correlations of gene essentiality scores across different screening platforms in human A375 cells. We consider all genes, including the non-essential gene controls in our screen, shared across all screens (n = 1,165). Gene-level score from our screen is calculated as the median of the top 4 common guides per junction and then top 4 junctions per gene. (D) Pearson correlation of LFC for ∼258 gRNAs shared between a previous HEK293FT screen (Wessels et al. 2023) and our A375 screen. Left plot compares Day 21 LFC between biological replicates of our A375 screen and the right plot compares the average Day 21 LFC of the two replicates in our A375 screen to the average Day 30 LFC Wessels et al screen in HEK293FT. (E) LFC for 1,992 non-essential (NE) gene (n = 413 genes) control common junction gRNAs versus TPM gene expression in A375. Only a very slight reduction in LFC is seen for high TPM genes, indicating that Cas13d collateral activity is not a major concern.

**Supplemental Figure 2.**
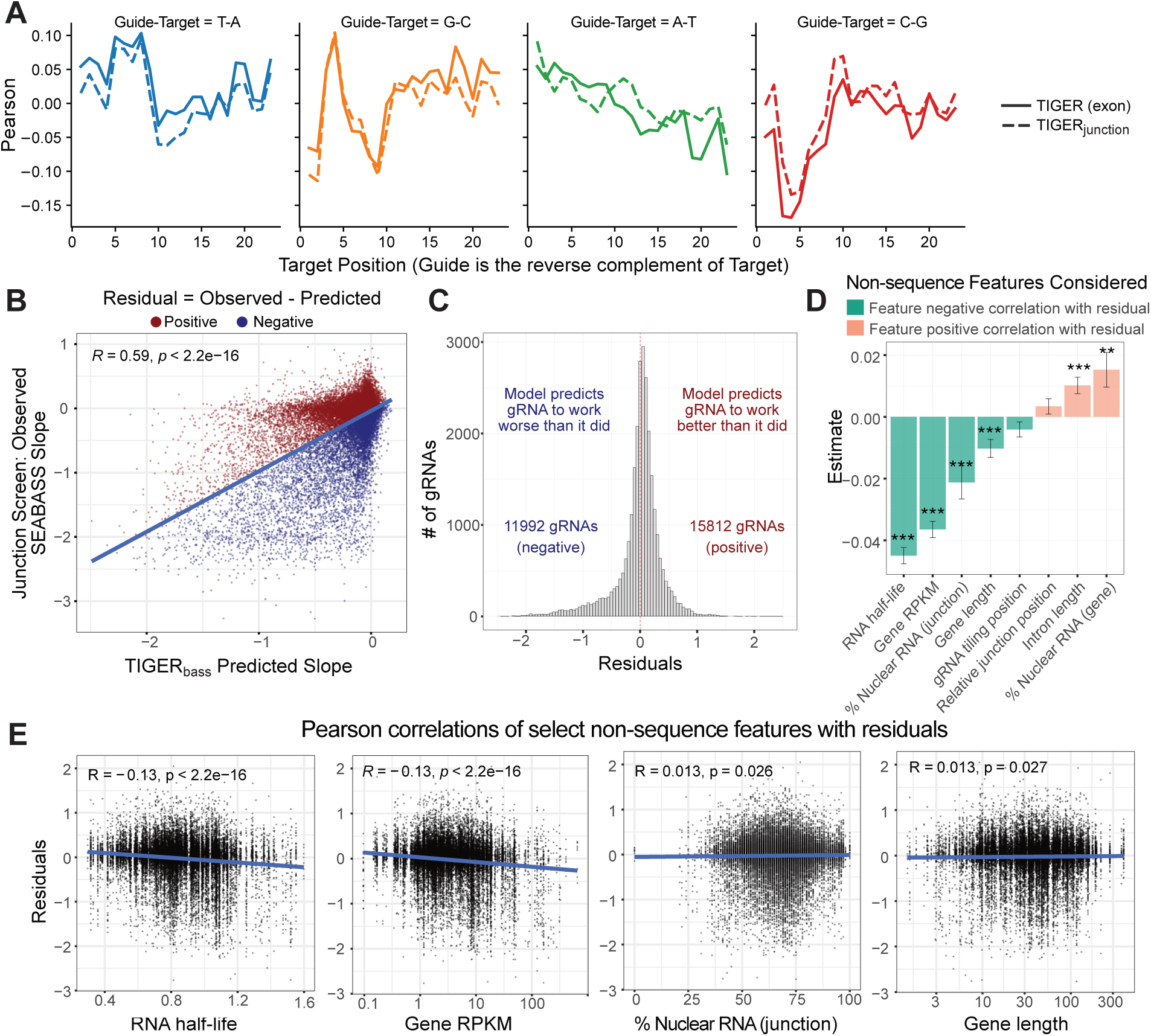
(A) Pearson plots comparing the correlation between positional nucleotide identity and LFC for TIGER and TIGERjunction. A negative correlation indicates that the presence of that nucleotide at that position results in a more active gRNA (i.e more negative LFC). (B) Pearson correlation of TIGERbass predicted slopes versus SEABASS observed slopes for junction targeting gRNAs in the essentiality screen. Residuals are calculated as observed minus predicted slope. Positive and negative residuals are colored red and blue respectively. (C) Residuals calculated in (B) shown as a histogram as the number of gRNAs/bin. Positive residuals indicate that model predictions outweighed observed gRNA efficacies in our screen. (D) Coefficient estimates from a multiple linear regression model for residuals using additional non-sequence features. P-values: * <0.05, ** <0.01 and *** <0.001. (E) Pearson correlations for residuals versus values from selected non-sequence features in (D).

**Supplemental Figure 3.**
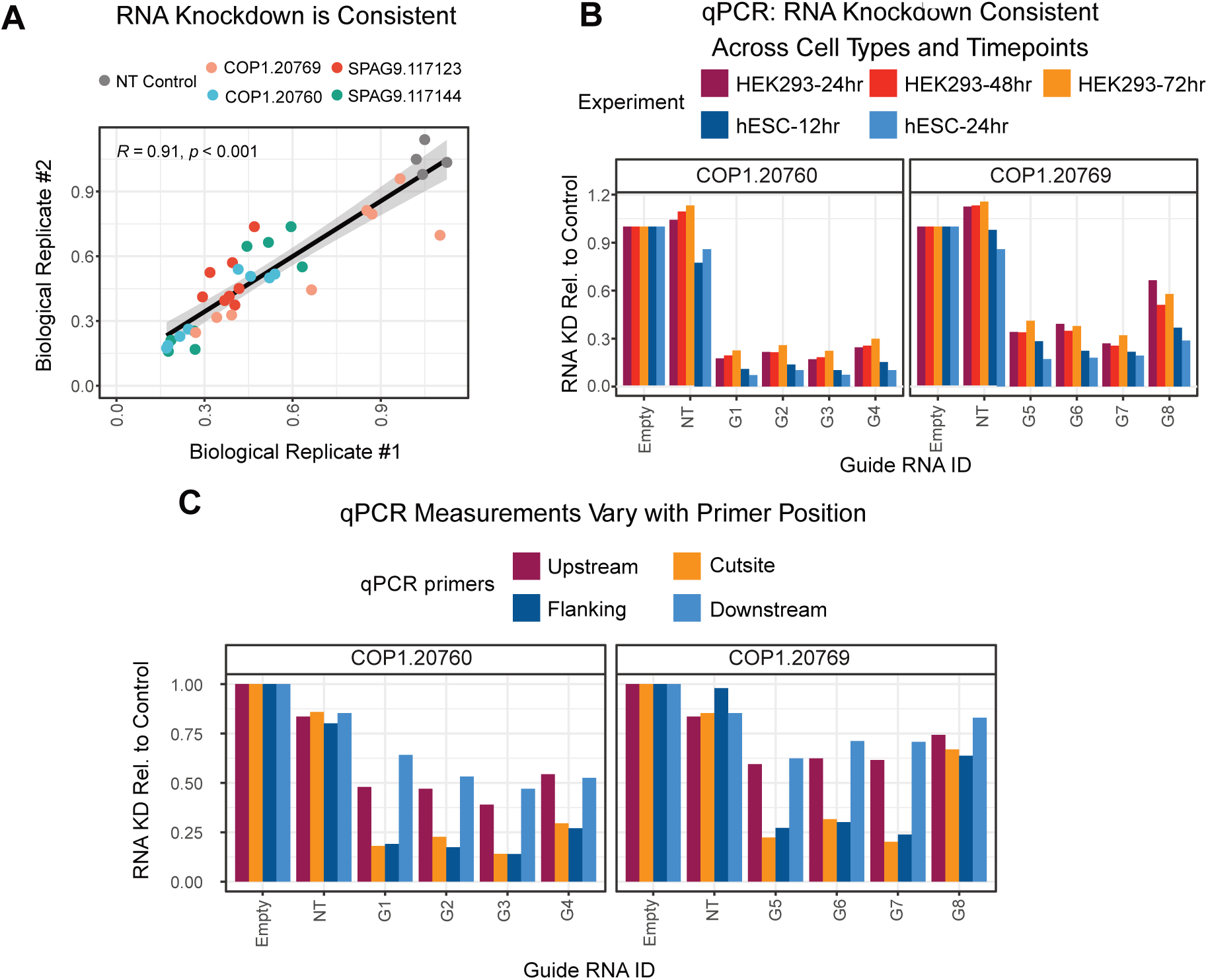
(A) Pearson correlation of RT-qPCR measurements between two biological replicates for *COP1* and *SPAG9*. Each point represents an average of 3 RT-qPCR technical replicates and is colored based on the junction. There are eight measurements per junction because two RT-qPCR assays were performed on each sample using two different primer sets. (B) Barplots show *COP1* RNA knockdown across two different cell types, HEK293 and hESC, and three timepoints of doxycycline-induced expression of Cas13. Each bar represents the average of 3 RT-qPCR technical replicates from a single biological replicate. (C) *COP1* RNA knockdown measured by RT-qPCR. All 4 bars for each gRNA ID (e.g. “G1”) on the x-axis represent measurements taken from the same sample but with different primers. Primers were designed both upstream of the gRNA cutsite (maroon), one primer overlapping the gRNA cutsite (gold), primers flanking the gRNA cutsite (navy blue), or both downstream of the gRNA cutsite (light blue). Each bar represents the average of 3 RT-qPCR technical replicates.

**Supplemental Figure 4.**
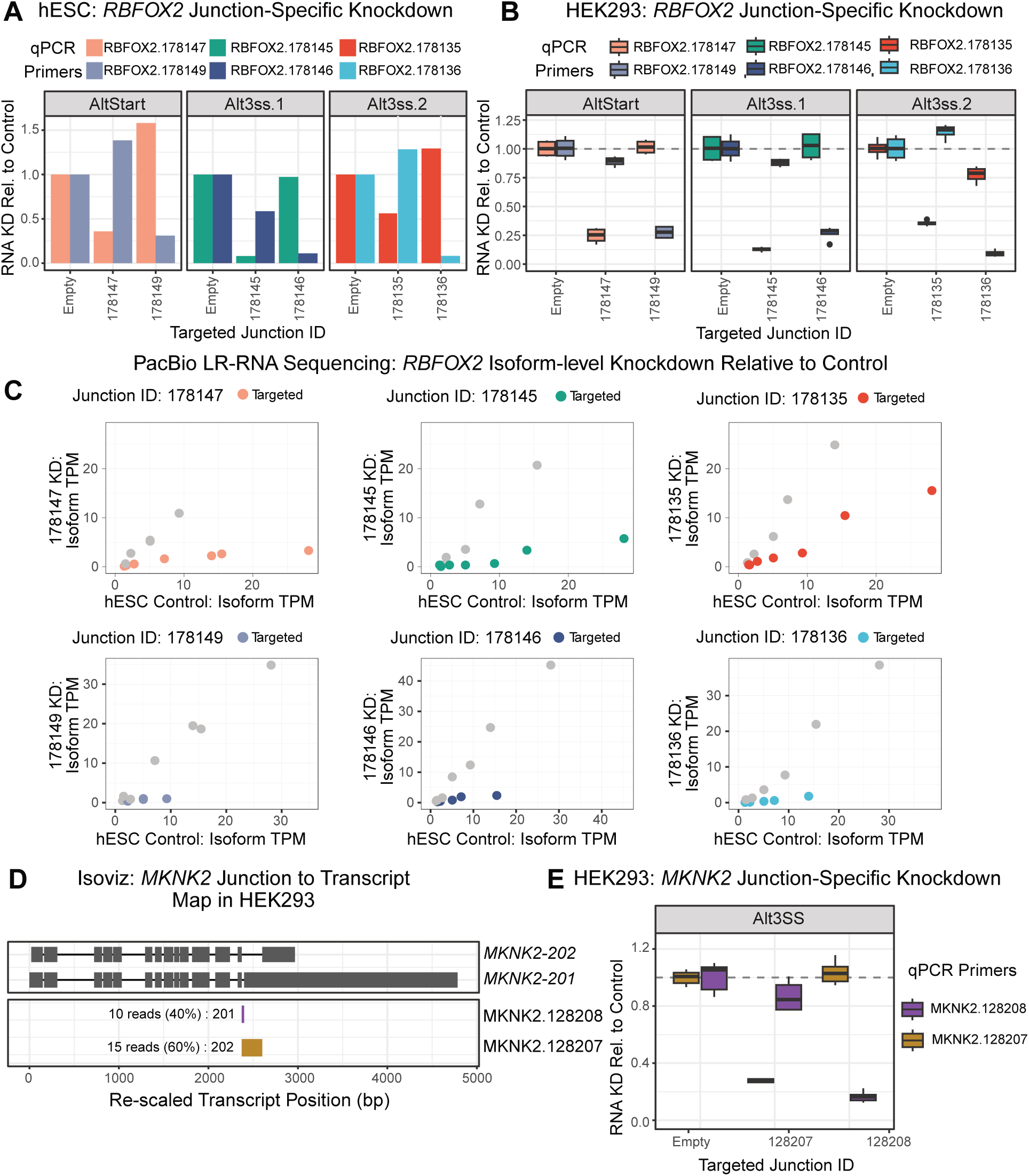
(A) *RBFOX2* junction-specific RNA knockdown in hESCs measured by RT-qPCR. RNA samples are the same ones used for the PacBio LRS in Figure 4. Bar plots represent an average of two RT-qPCR measurements from one biological replicate. (B) *RBFOX2* junction-specific RNA knockdown in HEK293 cells measured by RT-qPCR relative to cells transfected with empty guide cassette as a control. Junction IDs and boxplot colors correspond to junctions in Figure 4A. Each boxplot represents the average of 3 RT-qPCR measurements from two biological replicates. (C) Alternative view of the data plotted in Figure 4B-D. Transcript per million (TPM) values for each *RBFOX2* isoform is calculated from full length reads from PacBio LRS. TPM for all *RBFOX2* isoforms present in the hESC control (isoforms > 0.6 TPM or 10 full length reads) is compared to the TPM in each of six samples transfected with junction-specific gRNAs. Each point represents a transcript and is colored according to whether it contains the junction being targeted in that sample. All colors and IDs match those in Figure 4 and panels A and B here. (D) Isoviz output plotting two *MKNK2* transcripts that contain an alternative 3’ SS event at junctions 128208 (purple) and 128207 (gold). The text next to each intron corresponds to the number of reads overlapping that junction in HEK293 cells and the specific transcript that junction maps to. (E) *MKNK2* junction-specific RNA knockdown in HEK293 cells measured by RT-qPCR. Junction IDs and boxplot colors correspond to junctions in (D). Each boxplot represents two RT-qPCR measurements from two biological replicates.

## Supplemental Tables

**Supplemental Table S1: Oligonucleotides used in this paper.** Table gives all oligonucleotides used in this manuscript. This includes oligos used for cloning and library preparation. ‘Knowles Lab Primer Name’ is our internal primer id. HPLC-purified is given in parentheses for the subset of oligos that were special ordered; all other oligos were ordered with default conditions. ‘Paper Primer Name’ gives the id specific to this paper (if the id is used in figure legends), ‘Sequence’ is the oligo sequence, ‘Assay’ is either RT-qPCR or PCR. ‘Figure’ includes the specific figure(s) where the oligo was used or the methods section which mentions the oligo.

**Supplemental Table S2: gRNA sequences used in this paper.** Table gives all gRNAs used in this manuscript. ‘Guide RNA Sequence’ is the 23bp guide RNA sequence that is the reverse complement of the target RNA, without any added sequences for cloning. This would be considered the “top” sequence; for cloning, top and bottom sequences are ordered with added 5’ overhangs specific to the gRNA vector backbone. ‘Gene’ is the gene target of the gRNA. ‘Paper Guide ID’ gives the id that is used throughout this paper’s figure legends, while ‘Universal ID’ gives the unique id that we generated considering all possible junction gRNAs (8 guides per junction) in GENCODE v41 basic annotations. It combines the universal junction id with the tiling position within the junction (1-8). Importantly, RBFOX2 G11 position is outside the normal tiling position, so it does not have a universal ID. ‘Figure’ indicates the specific figure(s) where the gRNA was used.

**Supplemental Table S3: Sequencing data generated in this paper.** Table is divided into three sections based on category of sequencing: ‘Screen Libraries’, ‘RNA-seq Libraries’, and ‘LR-RNA-seq Libraries’ which correspond to GEO SubSeries GSE242106, GSE242105, and GSE242104, respectively. If a column is missing in one category versus the other, it means that column is not relevant to that data type. ‘ID’ gives the name of the dataset. ‘Date sequenced’ gives the date the data was obtained from sequencing. ‘Cell type’ gives the cell line the experiments were performed in. ‘Biological Replicate’ refers to independent experiments from different dates and ‘Technical Replicate’ refers to re-sequencing of the same library on multiple occasions. ‘Timepoint’ refers to the days or hours of dox-induction (which corresponds to Cas13d expression for all data here). ‘Treatment’ gives the dose of doxycycline added to the cells for the given experiment. ‘Stable Cell Line Type’ is either piggyBac or lentivirus. ‘ERCC spike-ins’ gives the amount of ERCC spike-ins added to RNA prior to library prep. See methods for more details. ‘Sequencing ID’, ‘Forward barcode’, and ‘Reverse barcode’ correspond to the primer IDs and sequences in Table S1 used for library prep. These are also the barcodes used for de-mulitplexing samples.

## Acknowledgements

We thank PacBio for partnering with us to generate data for this paper using a pre-commercial product. We thank Gloria Sheynkman and members of her lab for feedback on Isoviz and helpful discussions. M.D.S. was supported by NIH/NIGMS (F32GM142213). N.E.S. is supported by NYU and NYGC startup funds, NIH/NHGRI (DP2HG010099), NIH/NCI (R01CA279135, R21CA272345), NIH/NIAID (R01AI176601), the MacMillan Center for the Study of the Non-Coding Cancer Genome, and the Simons Foundation for Autism Research Initiative (896724). D.A.K. is supported by Columbia University and NYGC startup funds, and NIH/NCI (R21CA272345). A.S. and D.A.K. were supported by NSF CAREER DBI2146398.

## Author Contributions

M.D.S., L.P., and D.A.K. conceived the study. L.P. and H.W. performed the CRISPR screen. M.D.S., C.H., and S.H.P. performed all additional experiments. M.D.S., A.S., K.I., A.D., S.H.P., and D.A.K performed analyses. M.D.S. and K.I. developed Isoviz. A.D. and D.A.K developed SEABASS. A.S. developed the variants of TIGER. H.W. and N.E.S. provided reagents and advice. M.D.S and D.A.K. wrote the paper with input from co-authors. M.D.S. and D.A.K. supervised the study.

## Availability of data and materials

All sequencing data generated in this study has been deposited to NCBI Gene Expression Omnibus (GEO) under accession number GSE242107. SEABASS is publicly available at https://github.com/daklab/seabass and on pypi (https://pypi.org/project/seabass/0.0.5/). Isoviz R package is available at https://github.com/daklab/isoviz. TIGER is available at https://github.com/daklab/tiger with a user-friendly website for gRNA prediction at tiger.nygenome.org.

Once the paper is published, we plan to submit the plasmids generated in this study to Addgene.

## References

Abudayyeh, Omar O., Jonathan S. Gootenberg, Silvana Konermann, Julia Joung, Ian M. Slaymaker, David B. T. Cox, Sergey Shmakov, et al. 2016. “C2c2 Is a Single-Component Programmable RNA-Guided RNA-Targeting CRISPR Effector.” Science 353 (6299). 10.1126/science.aaf5573.

Agarwal, Vikram, and David R. Kelley. 2022. “The Genetic and Biochemical Determinants of mRNA Degradation Rates in Mammals.” Genome Biology 23 (1): 245.

Ai, Yuxi, Dongming Liang, and Jeremy E. Wilusz. 2022. “CRISPR/Cas13 Effectors Have Differing Extents of off-Target Effects That Limit Their Utility in Eukaryotic Cells.” Nucleic Acids Research 50 (11): e65.

Al’Khafaji, Aziz M., Jonathan T. Smith, Kiran V. Garimella, Mehrtash Babadi, Victoria Popic, Moshe Sade-Feldman, Michael Gatzen, et al. 2023. “High-Throughput RNA Isoform Sequencing Using Programmed cDNA Concatenation.” *Nature Biotechnology*, June, 1–5.

Baralle, Francisco E., and Jimena Giudice. 2017. “Alternative Splicing as a Regulator of Development and Tissue Identity.” Nature Reviews. Molecular Cell Biology 18 (7): 437–51.

Belluti, Silvia, Valentina Semeghini, Giovanna Rigillo, Mirko Ronzio, Daniela Benati, Federica Torricelli, Luca Reggiani Bonetti, et al. 2021. “Alternative Splicing of NF-YA Promotes Prostate Cancer Aggressiveness and Represents a New Molecular Marker for Clinical Stratification of Patients.” Journal of Experimental & Clinical Cancer Research: CR 40 (1): 362.

Burris, Brandon Joseph Davis, Adrian Moises Molina Vargas, Brandon J. Park, and Mitchell R. O’Connell. 2022. “Optimization of Specific RNA Knockdown in Mammalian Cells with CRISPR-Cas13.”

Cadiñanos, Juan, and Allan Bradley. 2007. “Generation of an Inducible and Optimized piggyBac Transposon System.” Nucleic Acids Research 35 (12): e87.

Celotto, Alicia M., and Brenton R. Graveley. 2002. “Exon-Specific RNAi: A Tool for Dissecting the Functional Relevance of Alternative Splicing.”

Cheng, Xiaolong, Zexu Li, Ruocheng Shan, Zihan Li, Shengnan Wang, Wenchang Zhao, Han Zhang, et al. 2023. “Modeling CRISPR-Cas13d on-Target and off-Target Effects Using Machine Learning Approaches.” Nature Communications 14 (1): 1–14.

Chen, Sidi, Neville E. Sanjana, Kaijie Zheng, Ophir Shalem, Kyungheon Lee, Xi Shi, David A. Scott, et al. 2015. “Genome-Wide CRISPR Screen in a Mouse Model of Tumor Growth and Metastasis.” Cell 160 (6): 1246–60.

Chen, Sisi, Salima Benbarche, and Omar Abdel-Wahab. 2021. “Splicing Factor Mutations in Hematologic Malignancies.” Blood 138 (8): 599–612.

Cotto, Kelsy C., Yang-Yang Feng, Avinash Ramu, Megan Richters, Sharon L. Freshour, Zachary L. Skidmore, Huiming Xia, et al. 2023. “Integrated Analysis of Genomic and Transcriptomic Data for the Discovery of Splice-Associated Variants in Cancer.” Nature Communications 14 (1): 1589.

Damianov, Andrey, and Douglas L. Black. 2010. “Autoregulation of Fox Protein Expression to Produce Dominant Negative Splicing Factors.”

Danan-Gotthold, Miri, Regina Golan-Gerstl, Eli Eisenberg, Keren Meir, Rotem Karni, and Erez Y. Levanon. 2015. “Identification of Recurrent Regulated Alternative Splicing Events across Human Solid Tumors.” Nucleic Acids Research 43 (10): 5130–44.

Davies, Rebecca, Ling Liu, Sheng Taotao, Natasha Tuano, Richa Chaturvedi, Kie Kyon Huang, Catherine Itman, et al. 2021. “CRISPRi Enables Isoform-Specific Loss-of-Function Screens and Identification of Gastric Cancer-Specific Isoform Dependencies.” Genome Biology 22 (1): 47.

Ding, Sheng, Xiaohui Wu, Gang Li, Min Han, Yuan Zhuang, and Tian Xu. 2005. “Efficient Transposition of the piggyBac (PB) Transposon in Mammalian Cells and Mice.” Cell 122 (3): 473–83.

Dobin, Alexander, Carrie A. Davis, Felix Schlesinger, Jorg Drenkow, Chris Zaleski, Sonali Jha, Philippe Batut, Mark Chaisson, and Thomas R. Gingeras. 2013. “STAR: Ultrafast Universal RNA-Seq Aligner.” Bioinformatics 29 (1): 15–21.

Doench, John G., Nicolo Fusi, Meagan Sullender, Mudra Hegde, Emma W. Vaimberg, Katherine F. Donovan, Ian Smith, et al. 2016. “Optimized sgRNA Design to Maximize Activity and Minimize off-Target Effects of CRISPR-Cas9.” Nature Biotechnology 34 (2): 184–91.

East-Seletsky, Alexandra, Mitchell R. O’Connell, Spencer C. Knight, David Burstein, Jamie H. D. Cate, Robert Tjian, and Jennifer A. Doudna. 2016. “Two Distinct RNase Activities of CRISPR-C2c2 Enable Guide-RNA Processing and RNA Detection.” Nature. 10.1038/nature19802.

Frankish, Adam, Mark Diekhans, Irwin Jungreis, Julien Lagarde, Jane E. Loveland, Jonathan M. Mudge, Cristina Sisu, et al. 2021. “GENCODE 2021.” Nucleic Acids Research 49 (D1): D916–23.

Gapinske, Michael, Alan Luu, Jackson Winter, Wendy S. Woods, Kurt A. Kostan, Nikhil Shiva, Jun S. Song, and Pablo Perez-Pinera. 2018. “CRISPR-SKIP: Programmable Gene Splicing with Single Base Editors.” Genome Biology 19 (1): 107.

Gonatopoulos-Pournatzis, Thomas, Michael Aregger, Kevin R. Brown, Shaghayegh Farhangmehr, Ulrich Braunschweig, Henry N. Ward, Kevin C. H. Ha, et al. 2020. “Genetic Interaction Mapping and Exon-Resolution Functional Genomics with a Hybrid Cas9-Cas12a Platform.” Nature Biotechnology 38 (5): 638–48.

GTEx Consortium. 2020. “The GTEx Consortium Atlas of Genetic Regulatory Effects across Human Tissues.” Science 369 (6509): 1318–30.

Gueroussov, Serge, Thomas Gonatopoulos-Pournatzis, Manuel Irimia, Bushra Raj, Zhen-Yuan Lin, Anne-Claude Gingras, and Benjamin J. Blencowe. 2015. “An Alternative Splicing Event Amplifies Evolutionary Differences between Vertebrates.” Science 349 (6250): 868–73.

Gusev, Alexander, Nicholas Mancuso, Hyejung Won, Maria Kousi, Hilary K. Finucane, Yakir Reshef, Lingyun Song, et al. 2018. “Transcriptome-Wide Association Study of Schizophrenia and Chromatin Activity Yields Mechanistic Disease Insights.” Nature Genetics 50 (4): 538–48.

Hir, Hervé Le, Jérôme Saulière, and Zhen Wang. 2015. “The Exon Junction Complex as a Node of Post-Transcriptional Networks.” Nature Reviews. Molecular Cell Biology 17 (1): 41–54.

Kahles, André, Kjong-Van Lehmann, Nora C. Toussaint, Matthias Hüser, Stefan G. Stark, Timo Sachsenberg, Oliver Stegle, et al. 2018. “Comprehensive Analysis of Alternative Splicing Across Tumors from 8,705 Patients.” Cancer Cell 34 (2): 211–24.e6.

Kirk, Jessime M., Susan O. Kim, Kaoru Inoue, Matthew J. Smola, David M. Lee, Megan D. Schertzer, Joshua S. Wooten, et al. 2018. “Functional Classification of Long Non-Coding RNAs by K-Mer Content.” Nature Genetics 50 (10): 1474–82.

Kushawah, Gopal, Luis Hernandez-Huertas, Joaquin Abugattas-Nuñez Del Prado, Juan R. Martinez-Morales, Michelle L. DeVore, Huzaifa Hassan, Ismael Moreno-Sanchez, et al. 2020. “CRISPR-Cas13d Induces Efficient mRNA Knockdown in Animal Embryos.” Developmental Cell 54 (6): 805–17.e7.

Langmead, Ben, Cole Trapnell, Mihai Pop, and Steven L. Salzberg. 2009. “Ultrafast and Memory-Efficient Alignment of Short DNA Sequences to the Human Genome.” Genome Biology 10 (3): R25.

Legut, Mateusz, Zoran Gajic, Maria Guarino, Zharko Daniloski, Jahan A. Rahman, Xinhe Xue, Congyi Lu, et al. 2022. “A Genome-Scale Screen for Synthetic Drivers of T Cell Proliferation.” Nature 603 (7902): 728–35.

Lewis, Benjamin P., Richard E. Green, and Steven E. Brenner. 2003. “Evidence for the Widespread Coupling of Alternative Splicing and Nonsense-Mediated mRNA Decay in Humans.” Proceedings of the National Academy of Sciences 100 (1): 189–92.

Liao, Yang, Gordon K. Smyth, and Wei Shi. 2013. “featureCounts: An Efficient General Purpose Program for Assigning Sequence Reads to Genomic Features.” Bioinformatics 30 (7): 923–30.

Li, Heng. 2018. “Minimap2: Pairwise Alignment for Nucleotide Sequences.” Bioinformatics 34 (18): 3094–3100.

Li, Heng, Bob Handsaker, Alec Wysoker, Tim Fennell, Jue Ruan, Nils Homer, Gabor Marth, Goncalo Abecasis, and Richard Durbin. 2009. “The Sequence Alignment/Map Format and SAMtools.” Bioinformatics 25 (16): 2078–79.

Li, Shenglan, Anqi Zhang, Haipeng Xue, Dali Li, and Ying Liu. 2017. “One-Step piggyBac Transposon-Based CRISPR/Cas9 Activation of Multiple Genes.” Molecular Therapy. Nucleic Acids 8 (September): 64–76.

Li, Yang I., Bryce van de Geijn, Anil Raj, David A. Knowles, Allegra A. Petti, David Golan, Yoav Gilad, et al. 2016. “RNA Splicing Is a Primary Link between Genetic Variation and Disease.” Science 352 (6285): 600–604.

Li, Yang I., David A. Knowles, Jack Humphrey, Alvaro N. Barbeira, Scott P. Dickinson, Hae Kyung Im, and Jonathan K. Pritchard. 2017. “Annotation-Free Quantification of RNA Splicing Using LeafCutter.” Nature Genetics 50 (1): 151–58.

Love, Michael I., Wolfgang Huber, and Simon Anders. 2014. “Moderated Estimation of Fold Change and Dispersion for RNA-Seq Data with DESeq2.” Genome Biology 15 (12): 550.

Lundberg, Scott M., and Su-In Lee. 2017. “A Unified Approach to Interpreting Model Predictions.” Advances in Neural Information Processing Systems 30.

Maimon, Avraham, Maxim Mogilevsky, Asaf Shilo, Regina Golan-Gerstl, Akram Obiedat, Vered Ben-Hur, Ilana Lebenthal-Loinger, et al. 2014. “Mnk2 Alternative Splicing Modulates the p38-MAPK Pathway and Impacts Ras-Induced Transformation.” Cell Reports 7 (2): 501–13.

Martin, Marcel. 2011. “Cutadapt Removes Adapter Sequences from High-Throughput Sequencing Reads.” EMBnet.journal 17 (1): 10–12.

Miller, Rachel M., Ben T. Jordan, Madison M. Mehlferber, Erin D. Jeffery, Christina Chatzipantsiou, Simi Kaur, Robert J. Millikin, et al. 2022. “Enhanced Protein Isoform Characterization through Long-Read Proteogenomics.” Genome Biology 23 (1): 69.

Morgens, David W., Richard M. Deans, Amy Li, and Michael C. Bassik. 2016. “Systematic Comparison of CRISPR/Cas9 and RNAi Screens for Essential Genes.” Nature Biotechnology 34 (6): 634–36.

Pan, Qun, Ofer Shai, Leo J. Lee, Brendan J. Frey, and Benjamin J. Blencowe. 2008. “Deep Surveying of Alternative Splicing Complexity in the Human Transcriptome by High-Throughput Sequencing.” Nature Genetics 40 (12): 1413–15.

Prinos, Panagiotis, Daniel Garneau, Jean-François Lucier, Daniel Gendron, Sonia Couture, Marianne Boivin, Jean-Philippe Brosseau, et al. 2011. “Alternative Splicing of SYK Regulates Mitosis and Cell Survival.” Nature Structural & Molecular Biology 18 (6): 673–79.

Raj, Towfique, Yang I. Li, Garrett Wong, Jack Humphrey, Minghui Wang, Satesh Ramdhani, Ying-Chih Wang, et al. 2018. “Integrative Transcriptome Analyses of the Aging Brain Implicate Altered Splicing in Alzheimer’s Disease Susceptibility.” Nature Genetics 50 (11): 1584–92.

Reese, Fairlie, Brian Williams, Gabriela Balderrama-Gutierrez, Dana Wyman, Muhammed Hasan Çelik, Elisabeth Rebboah, Narges Rezaie, et al. 2023. “The ENCODE4 Long-Read RNA-Seq Collection Reveals Distinct Classes of Transcript Structure Diversity.” bioRxiv. https://www.ncbi.nlm.nih.gov/pmc/articles/PMC10245583.

Regan-Fendt, Kelly E., Jielin Xu, Mallory DiVincenzo, Megan C. Duggan, Reena Shakya, Ryejung Na, William E. Carson 3rd, Philip R. O. Payne, and Fuhai Li. 2019. “Synergy from Gene Expression and Network Mining (SynGeNet) Method Predicts Synergistic Drug Combinations for Diverse Melanoma Genomic Subtypes.” NPJ Systems Biology and Applications 5 (February): 6.

Reixachs-Solé, Marina, and Eduardo Eyras. 2022. “Uncovering the Impacts of Alternative Splicing on the Proteome with Current Omics Techniques.” Wiley Interdisciplinary Reviews. RNA 13 (4): e1707.

Rogalska, Malgorzata Ewa, Claudia Vivori, and Juan Valcárcel. 2023. “Regulation of Pre-mRNA Splicing: Roles in Physiology and Disease, and Therapeutic Prospects.” Nature Reviews. Genetics 24 (4): 251–69.

Sanson, Kendall R., Ruth E. Hanna, Mudra Hegde, Katherine F. Donovan, Christine Strand, Meagan E. Sullender, Emma W. Vaimberg, et al. 2018. “Optimized Libraries for CRISPR-Cas9 Genetic Screens with Multiple Modalities.” Nature Communications 9 (1): 1–15.

Schertzer, Megan D., Eliza Thulson, Keean C. A. Braceros, David M. Lee, Emma R. Hinkle, Ryan M. Murphy, Susan O. Kim, Eva C. M. Vitucci, and J. Mauro Calabrese. 2019. “A piggyBac-Based Toolkit for Inducible Genome Editing in Mammalian Cells.” RNA 25 (8): 1047–58.

Scotti, Marina M., and Maurice S. Swanson. 2015. “RNA Mis-Splicing in Disease.” Nature Reviews. Genetics 17 (1): 19–32.

Seiler, Michael, Shouyong Peng, Anant A. Agrawal, James Palacino, Teng Teng, Ping Zhu, Peter G. Smith, Cancer Genome Atlas Research Network, Silvia Buonamici, and Lihua Yu. 2018. “Somatic Mutational Landscape of Splicing Factor Genes and Their Functional Consequences across 33 Cancer Types.” Cell Reports 23 (1): 282–96.e4.

Shalem, Ophir, Neville E. Sanjana, Ella Hartenian, Xi Shi, David A. Scott, Tarjei Mikkelson, Dirk Heckl, et al. 2014. “Genome-Scale CRISPR-Cas9 Knockout Screening in Human Cells.” Science 343 (6166): 84–87.

Shi, Peiguo, Michael R. Murphy, Alexis O. Aparicio, Jordan S. Kesner, Zhou Fang, Ziheng Chen, Aditi Trehan, Yang Guo, and Xuebing Wu. 2023. “Collateral Activity of the CRISPR/RfxCas13d System in Human Cells.” Communications Biology 6 (1): 1–8.

Stanley, Robert F., and Omar Abdel-Wahab. 2022. “Dysregulation and Therapeutic Targeting of RNA Splicing in Cancer.” Nature Cancer 3 (5): 536–46.

Tang, Alison D., Cameron M. Soulette, Marijke J. van Baren, Kevyn Hart, Eva Hrabeta-Robinson, Catherine J. Wu, and Angela N. Brooks. 2020. “Full-Length Transcript Characterization of SF3B1 Mutation in Chronic Lymphocytic Leukemia Reveals Downregulation of Retained Introns.” Nature Communications 11 (1): 1–12.

Thomas, James D., Jacob T. Polaski, Qing Feng, Emma J. De Neef, Emma R. Hoppe, Maria V. McSharry, Joseph Pangallo, et al. 2020. “RNA Isoform Screens Uncover the Essentiality and Tumor-Suppressor Activity of Ultraconserved Poison Exons.” Nature Genetics 52 (1): 84–94.

Tsherniak, Aviad, Francisca Vazquez, Phil G. Montgomery, Barbara A. Weir, Gregory Kryukov, Glenn S. Cowley, Stanley Gill, et al. 2017. “Defining a Cancer Dependency Map.” Cell 170 (3): 564–76.e16.

Villemaire, Jonathan, Isabelle Dion, Sherif Abou Elela, and Benoit Chabot. 2003. “Reprogramming Alternative Pre-Messenger RNA Splicing through the Use of Protein-Binding Antisense Oligonucleotides.” The Journal of Biological Chemistry 278 (50): 50031– 39.

Vuong, Celine K., Douglas L. Black, and Sika Zheng. 2016. “The Neurogenetics of Alternative Splicing.” Nature Reviews. Neuroscience 17 (5): 265–81.

Wang, Eric T., Rickard Sandberg, Shujun Luo, Irina Khrebtukova, Lu Zhang, Christine Mayr, Stephen F. Kingsmore, Gary P. Schroth, and Christopher B. Burge. 2008. “Alternative Isoform Regulation in Human Tissue Transcriptomes.” Nature 456 (7221): 470–76.

Wang, Gang, Luhan Yang, Dennis Grishin, Xavier Rios, Lillian Y. Ye, Yong Hu, Kai Li, Donghui Zhang, George M. Church, and William T. Pu. 2017. “Efficient, Footprint-Free Human iPSC Genome Editing by Consolidation of Cas9/CRISPR and piggyBac Technologies.” Nature Protocols 12 (1): 88–103.

Wang, Qixue, Xing Liu, Junhu Zhou, Chao Yang, Guangxiu Wang, Yanli Tan, Ye Wu, Sijing Zhang, Kaikai Yi, and Chunsheng Kang. 2019. “The CRISPR-Cas13a Gene-Editing System Induces Collateral Cleavage of RNA in Glioma Cells.” Advancement of Science 6 (20): 1901299.

Wei, Jingyi, Peter Lotfy, Kian Faizi, Hugo Kitano, Patrick D. Hsu, and Silvana Konermann. 2021. “Deep Learning of Cas13 Guide Activity from High-Throughput Gene Essentiality Screening.” bioRxiv.

Wessels, Hans-Hermann, Alejandro Méndez-Mancilla, Xinyi Guo, Mateusz Legut, Zharko Daniloski, and Neville E. Sanjana. 2020. “Massively Parallel Cas13 Screens Reveal Principles for Guide RNA Design.” Nature Biotechnology 38 (6): 722–27.

Wessels, Hans-Hermann, Andrew Stirn, Alejandro Méndez-Mancilla, Eric J. Kim, Sydney K. Hart, David A. Knowles, and Neville E. Sanjana. 2023. “Prediction of on-Target and off-Target Activity of CRISPR-Cas13d Guide RNAs Using Deep Learning.” Nature Biotechnology, July. https://www.ncbi.nlm.nih.gov/pubmed/37400521.

Wilson, Matthew H., Craig J. Coates, and Alfred L. George Jr. 2007. “PiggyBac Transposon-Mediated Gene Transfer in Human Cells.” Molecular Therapy: The Journal of the American Society of Gene Therapy 15 (1): 139–45.

Yan, Qinghong, Sebastien M. Weyn-Vanhentenryck, Jie Wu, Steven A. Sloan, Ye Zhang, Kenian Chen, Jia Qian Wu, Ben A. Barres, and Chaolin Zhang. 2015. “Systematic Discovery of Regulated and Conserved Alternative Exons in the Mammalian Brain Reveals NMD Modulating Chromatin Regulators.” Proceedings of the National Academy of Sciences 112 (11): 3445–50.

Zhang, Yang, Tuan M. Nguyen, Xiao-Ou Zhang, Limei Wang, Tin Phan, John G. Clohessy, and Pier Paolo Pandolfi. 2021. “Optimized RNA-Targeting CRISPR/Cas13d Technology Outperforms shRNA in Identifying Functional circRNAs.” Genome Biology 22 (1): 41.

